# Data-driven large-scale genomic analysis reveals an intricate phylogenetic and functional landscape in J-domain proteins

**DOI:** 10.1101/2022.02.28.482344

**Authors:** Duccio Malinverni, Stefano Zamuner, Mathieu E. Rebeaud, Alessandro Barducci, Nadinath B. Nillegoda, Paolo De Los Rios

## Abstract

The 70 kDalton Heat shock protein (Hsp70) chaperone system is emerging as a central hub of the proteostasis network that helps maintain protein homeostasis in all organisms. The recruitment of Hsp70 to perform a vast array of different cellular functions is regulated by a family of co-chaperones known as J-domain proteins (JDP) that bear a small namesake J-domain, which is required to interact and drive the ATPase cycle of Hsp70s. Both prokaryotic and eukaryotic JDPs display staggering diversity in domain architecture (besides the ubiquitous J-domain), function, and cell localization. On the contrary, a relatively small number of Hsp70 paralogs exist in cells, suggesting a high degree of specificity, but also promiscuity, in the partnering between JDPs and Hsp70s. Very little is known about the JDP family, despite their essential role in cellular proteostasis, development, and the link to a broad range of human diseases. The number of JDP gene sequences identified across all kingdoms as a consequence of advancements in sequencing technology has exponentially increased, where it is now beyond the ability of careful manual curation. In this work, we first provide a broad overview of the JDP repertoire accessible from public databases, and then we use an automated classification scheme, based on Artificial Neural Networks (ANNs), to demonstrate that the sequences of J-domains carry sufficient discriminatory information to recover with high reliability the phylogeny, localization, and domain composition of the corresponding full-length JDP. By harnessing the interpretability of the ANNs, we find that many of the discriminatory sequence positions match to residues that form the interaction interface between the J-domain and Hsp70. This reveals that key residues within the J-domains have coevolved with their obligatory Hsp70 partners to build chaperone circuits for specific functions in cells.

## Introduction

Hsp70 is an essential and highly ubiquitous chaperone involved in a wide variety of constitutive and stress-related cellular processes, including the promotion of protein folding^1,2^, the prevention and reversal of the formation of cytotoxic aggregates^3–6^, the import of polypeptides into cell compartments ^7,8^, driving the (dis)assembly of oligomeric protein complexes^9–11^, the regulation of protein activity^12^ and the targeting of terminally damaged proteins for degradation^13–15^. Underlying these diverse functions is the fundamental ability of Hsp70 to bind substrate proteins upon a conformational change that depends on the nature of the bound nucleotide, which alternates between ATP and ADP. This biochemical cycle is stringently regulated by J-domain containing proteins (JDPs) that, together with the substrate, accelerate ATP hydrolysis in Hsp70s by orders of magnitude^16,17^, which results in ultra-affinity binding, namely a significant, non-equilibrium enhancement of the Hsp70 affinity for its substrates^18^. Subsequently, Hsp70 nucleotide exchange factors (NEFs) bind and accelerate ADP release, thus allowing rebinding of ATP and client protein release^19^ (Fig.1A).

**Figure 1.**
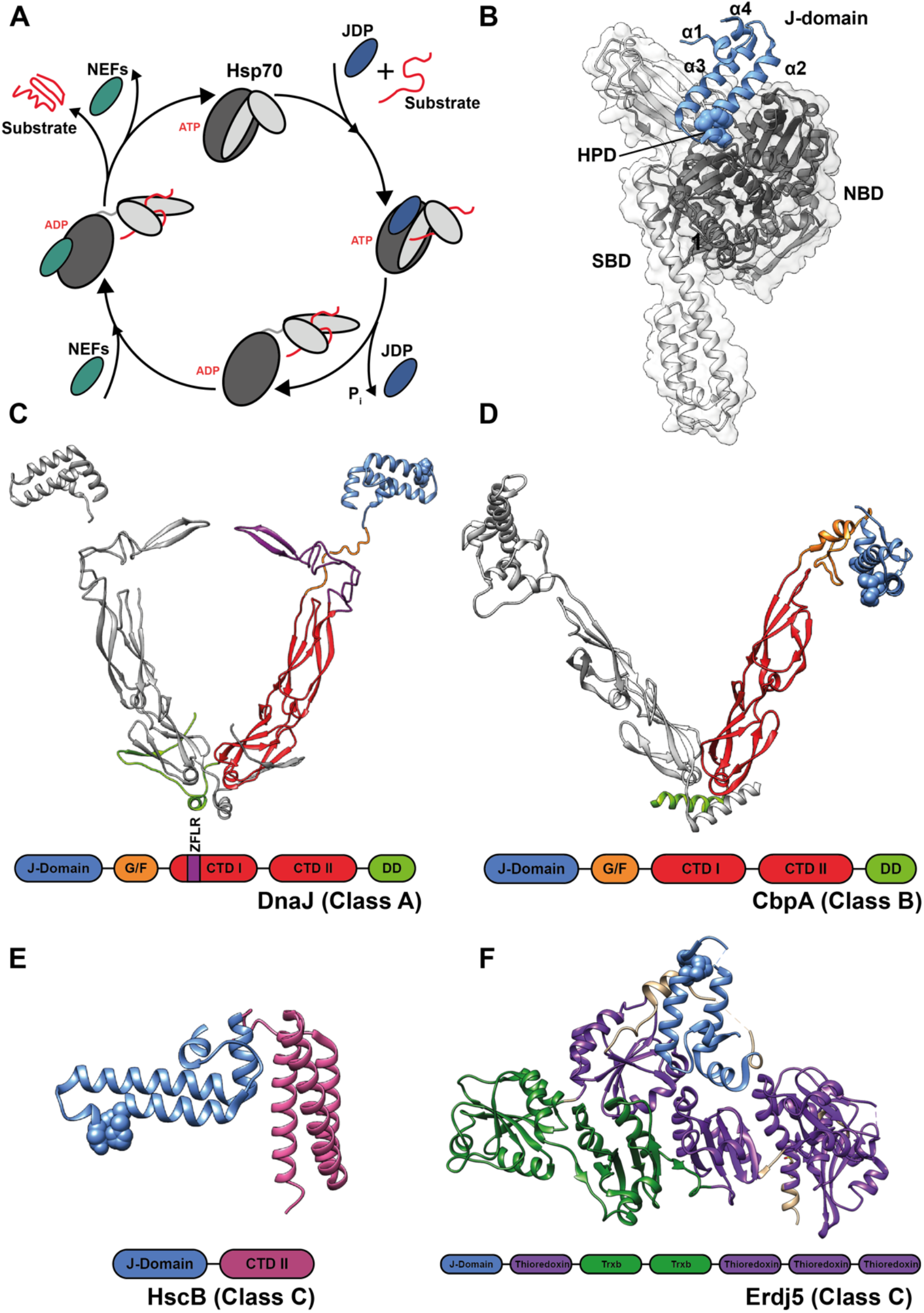
Structural views of the J-domain and domain architectures of JDPs. **A)** Schematic view of the canonical Hsp70 cycle. **B)** Structural view of the Hsp70-J-domain complex (PDB ID: 5NRO). NBD denotes the Hsp70 Nucleotide Binding Domain, SBD the Hsp70 Substrate Binding Domain. The four helices (α1-α4) forming the J-domain and the characteristic HPD motif are highlighted. **C)** Cytosolic class A JDP of *S. cerevisiae* (Ydj1) in its constitutive homo-dimeric form (the structure is a combination of the separate J-domain, PDB ID 5VSO, CTDs PDB ID 1NLT, and dimerization domains, PDB ID 1XAO); in lack of a known experimentally determined structure, the G/F rich linker between the J-domain and the first C-terminal domain has been hand-drawn on one of the two protomers. The different domains have been colored only on one protomer. The general architecture of class A JDPs is highlighted below the structure (G/F: glycine/phenylalanine rich linker; CTD: C-terminal, substrate binding domain; ZFLR: cysteine-rich, Zinc-Finger-Like Region; DD: dimerization domain). **C)** Class B JDP of *Thermus thermophilus* in its constitutive homo-dimeric form (PDB ID 1C3G). The different domains have been colored only on one protomer. The general architecture of class B JDPs is highlighted below the structure. **D)** *Escherichia coli* HscB (class C; PDB ID 1FPO) with its architecture (HSCB-C: C-terminal domain of HscB). **E)** *Mus musculus* Erdj5 (class C; PDB ID 5AYK) and its architecture.

All members of the JDP family invariably contain a J-domain, the namesake domain which is ∼63 residue-long. The J-domain is formed by a well-defined arrangement of four alpha helices (Fig.1B) with a highly-conserved Histidine-Proline-Aspartate (HPD) motif between helices 2 and 3. The helix II-HPD-helix III motif forms the primary interface for the interaction and stimulation of ATP hydrolysis in Hsp70^16,17,20–23.^ The remaining subdomains help target JDPs to specific locations, client proteins, or protein conformations (*e*.*g*. unfolded, misfolded, or aggregated polypeptides), thereby recruiting the action of Hsp70s to selected substrates^24,25^.

A classification scheme for JDPs has been suggested based on the overall domain architecture, resulting in three main classes^24,26–28^. Class A JDPs share the domain architecture of the prototypical DnaJ member of *Escherichia coli*. Members of this class consist of an N-terminal J-domain, followed by a glycine and phenylalanine (G/F) rich region and two successive C-terminal β-sandwich domains (CTDs), with a zinc-finger-like region (ZFLR) domain protruding from the first CTD, and are terminated by a short dimerization domain at the C-terminal (Fig.1C). The presence of an N-terminal J-domain fused with a G/F region was used as the primary domain signature to segregate some of the remaining JDPs to form the so-called class B. This relatively loose classification step, probably stemming from limited availability of JDP sequences at the conception of this class, comprises both the “canonical” type members that share the same domain architecture of class A with the exception of missing the ZFLR (Fig.1D), alongside more diverse ones, that are characterized by an N-terminal J-domain followed by a G/F rich region-only^27^. The G/F-rich region of both classes is typically considered largely unstructured. All the JDPs belonging to class A and some of the ones belonging to class B (also known collectively as Hsp40s owing to their approximate 40 kDa molecular mass) constitutively form homo-dimers through the C-terminal dimerization domain (Fig.1C and D), and bind a large variety of unfolded, misfolded, or aggregated substrates through one or multiple interaction sites located on their CTDs and, possibly for class A, on the ZFLR^29,30^. Thus, class A and B JDPs help recruit Hsp70s to their misfolded/aggregated substrates and contribute to the crowding of Hsp70s on the substrate^4^. In contrast, class C JDPs display a high variability in domain architectures (two examples are shown in Fig.1E and F) and have been historically classified by what they are not, namely neither A nor B, rather than by what they are. Indeed, while members of class C represent the vast majority of JDPs, and intervene in a wide range of unrelated biological pathways (*e*.*g*. auxilin, which is involved in clathrin cage disassembly; Hsc20, which regulates FeS cluster biogenesis; Sec63, which promotes post-translational protein import in the ER; Zuotin, which assists protein translation), they share with each other little or no common functional or phylogenetic features, except for the presence of a J-domain.

Here, we leveraged the recent exponential growth in genomic data and the improvements in homology detection tools to explore the relations between JDP sequences and their phylogenetic and functional classification at various resolutions. In particular, we focus on class C JDPs, and uncover several subclasses based on most common domain architectures. We then used Artificial Neural Networks (ANNs) to show that even the relatively short and simple sequences of the J-domains generally carry sufficient phylogenetic and functional signatures that may modulate JDP-Hsp70 chaperone circuits *in vivo*. These unique signatures reliably identify the architecture of the corresponding full-length JDP, its cell location, and the taxonomic clade. These findings allowed us to use amino acid sequence variations embedded in the J-domain to develop an ANN-based automated solution to curate existing and new JDP sequences reported in various organisms with high precision. By further identifying the residues that are most important for classification, we pinpointed the structural positions where the class- and organism-specific signatures are located.

From a more general perspective, we highlight how the combination of interpretable large-scale sequence analysis techniques and coarse-grained annotations can unlock the wealth of information that would otherwise remain hidden in exponentially growing, and thus manually unmanageable, sequence databases.

## Results

### Construction of an extensive annotated J-domain protein sequence dataset

The number of deposited JDP sequences in the UniProt repository^31^ has been exponentially growing over the past two decades, increasing by an order of magnitude every 6-7 years (Fig.2A), ranging from 10^3^ in year 2003 to 10^5^ in year 2018, and projected to reach 10^6^ sequences in years 2024-2025 and 10^7^ in the early 2030s. Using stringent homology search criteria, we extracted a total of 223’124 JDP sequences from existing genome databases (see Methods). The associated annotations allowed us to assign all JDPs to different phylogenetic classes at various taxonomic levels. Using sequence domain models from the Pfam database^32^, we further characterized the domain composition of full-length sequences, assigning to each JDP a tag describing the domain architecture of the protein. Protein-localization prediction algorithms were used to infer the most probable organelle of residence of eukaryotic JDPs (see Methods). This compound procedure allowed us to assign each JDP sequence in the dataset a set of *ground-truth* attributes comprising phylogenetic, architectural, and cellular localization features. In this context, the characteristics that describe, for instance, human DNAJA1 are: a) sequence composition, b) class (e.g. class A JDP based on the full-length domain composition), c) phylogeny (e.g. eukaryotic/metazoan/mammalian protein), and d) cell location (e.g. predominantly cytoplasmic).

**Figure 2.**
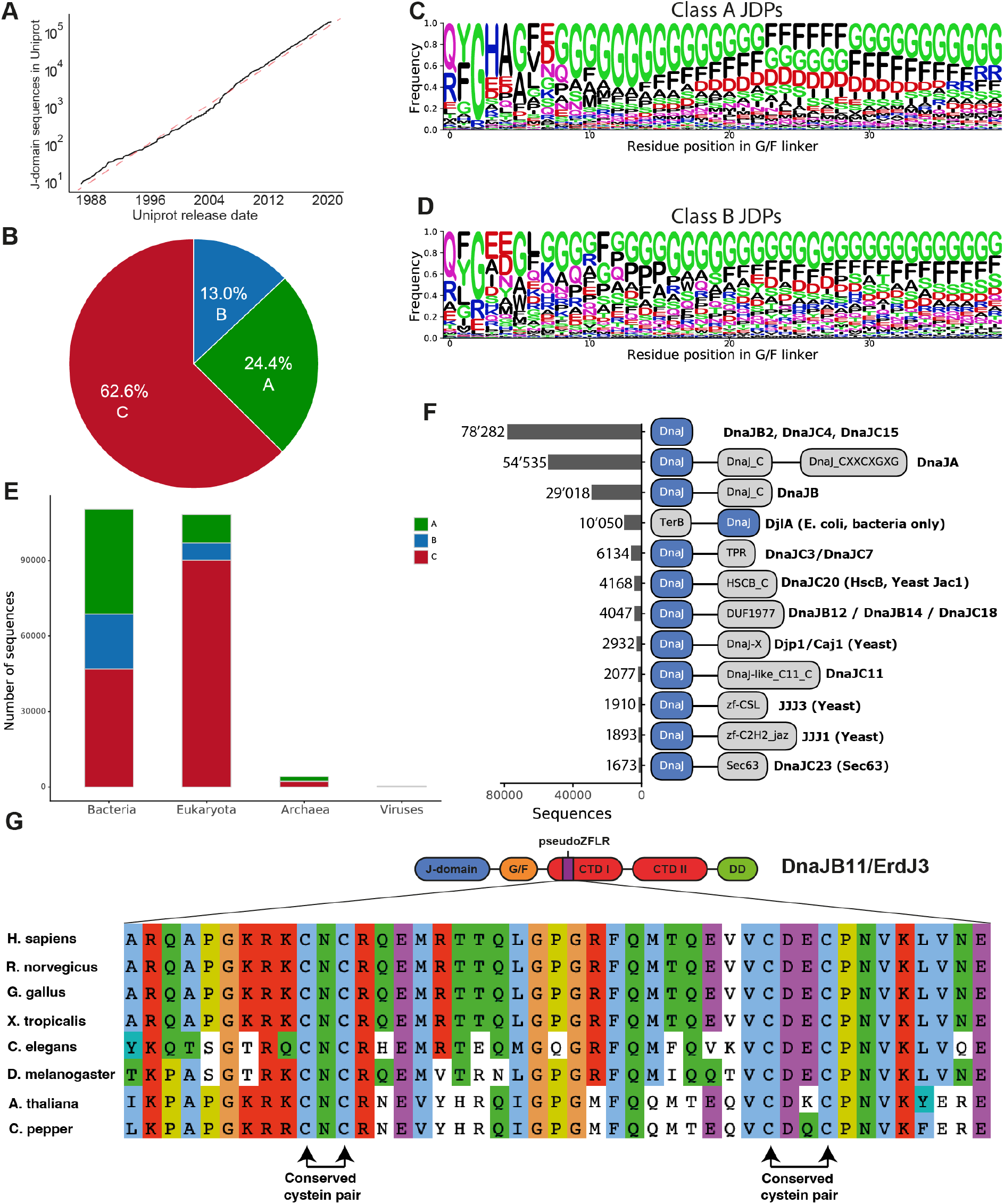
JDP dataset characterization. **A)** Growth of the number of JDP sequences found in the Uniprot Database over the last 3 decades. The red dashed line shows an exponential fit. **B)** Class proportions in the dataset. **C)** Amino-acid frequencies of the G/F linker region of class A JDPs. **D)** Amino-acid frequencies of the G/F linker region of class B JDPs. **E)** Number of JDPs in the different kingdoms, highlighted by their class (A: green; B: blue; C: red). **F)** The 12 most abundant J-protein domain architectures. Numbers on the left denote the number of sequences with this architecture in our dataset, and, on the right, we report the corresponding human, yeast, plant or bacterial proteins. **G)** Highlight of the evolutionary conservation of the cysteine-rich proto-ZFLR in ER-resident JDPs in 6 metazoa and 2 plants.

The correct identification and proper functional assignment of such a plethora of JDPs is incompatible with any careful manual curation. Instead, the rapidly growing size of the JDP repertoire will require rigorous, but automated characterization and analysis approaches trained on extensive datasets endowed with some reliable ground truth, defined according to different, complementary criteria. We provide such a dataset containing a comprehensive collection of richly annotated repertoire of JDPs in this work. This collection allowed us to investigate the ability of automated approaches to extract and rationalize the information it contains, while gaining new insights into this complex cochaperone family.

### Overall characterization of the diversity of the J-domain protein dataset

Based on the overall domain composition of JDPs, the three established functional classes were clearly identifiable: Class A and class B (members with the canonical Hsp40 domain architecture, Figs.1C and D) represented 24.4% and 13% of the dataset, respectively (Fig.2B), with the remaining JDPs (62.6%, corresponding to ∼140’000 sequences) assigned to class C. Approximately half of them (78’282) have a domain-architecture consisting of, according to the Pfam-based annotation, only the J-domain (Fig.2F). In some cases, this is due to the absence of any additional domains in the protein, while in others, it could be due to the inability of the Markov models used to effectively recognize variations of existing or potentially new domains. This is likely the case for DNAJB2, which has two ubiquitin binding modules (UIMs)^33,34^ that are nonetheless not detected using the Pfam Markov models. These observations clearly require curation not at the level of amino acid sequences, but at the level of the accuracy/coverage of the models. JDPs annotated with a single J-domain were slightly enriched in eukaryotic organisms (Supplementary Fig.1A) but found uniformly throughout all predicted subcellular locations (Supplementary Fig.1B). The full protein length distribution for JDPs annotated with a single J-domain was significantly shorter than for (annotated) JDPs consisting of multi-domain architectures (Supplementary Fig.1E). However, we also found a significant number of JDPs annotated with a single J-domain, but with relatively long protein sequences, which highlights the finite accuracy of automated domain annotation procedure.

Collectively, the 12 most populated domain architectures (including class A and B JDPs) comprised approximately 90% of the total dataset (196’719 sequences). Visual inspection of class A and B JDP sequences showed that beyond the clear difference due to the presence of the ZFLR, we could also distinguish sequences belonging to the two classes by the composition of their G/F-rich regions. Here, we show that the amino acid profiles of residues in the G/F regions of classes A and B are remarkably different, with the G/F-rich regions of class A showing a more marked segregation of phenylalanine, arginine, glycine and glutamate residues compared to those of class B JDPs (Fig.2C and D). This was quantitatively confirmed by a simple linear classification model which showed that the amino acid sequences (unaligned) of the G/F region alone were able to successfully discriminate JDPs of class A and B (classification accuracy of ∼95%, see FigS7 and Methods for details). These significant differences in amino acid compositions suggest that at least parts of the G/F rich region could have different functions beyond being a simple flexible linker. Recent observations show that in some class B members, a small helix in the G/F region docks with the J-domain, likely with functional consequences^35–38^. Furthermore, it was shown that some JDPs could, to a certain degree, undergo liquid-liquid phase separation with the help of the G/F regions^39^.

In agreement with previous observations^25^, class C members dominated the dataset (Fig.2B) mainly due to their overwhelming abundance in eukaryotes (Fig.2E and Supplementary Fig.2A for a more detailed analysis of class C abundance in the dataset). There were 1945 different architectures present in class C, which were built from 1558 distinct domains. Most of these domains were often classified as Domain of Unknown Function (DUFs) and showed little to no co-occurrence with the most common domains (Supplementary Fig.3). The only notable exceptions are JDPs containing multiple repeats of tetratricopeptides (TPR) associated with several different Pfam families (in Fig.2F and Supplementary Fig.3 all these TPR domains are collected in a single group). Inspection of the architectures of the most frequently observed JDP domains within class C revealed a very disparate phylogenetic makeup (Fig.2F). For instance, JDPs of the bacterial (*E. coli*) DjlA family, which formed the most abundant class C-subtype architecture, are present only in bacteria. The HscB family (human DNAJC20, Fig.1F), which participates in iron-sulfur cluster assembly pathway, is present in both prokaryotes and eukaryotes. Others, such as the yeast Sec63 (DNAJC18) localized in the endoplasmic reticulum (ER), (which shares almost the same domain composition as DNAJB12 and DNAJB14), DNAJC11 (DUF3995), and DNAJC3/ DNAJC7 were found only in eukaryotes. Some highly represented architectures obeyed an even finer phylogenetic distribution, with Djp1/Caj1, JJJ1, and JJJ3 (yeast names) present only in yeast (*e*.*g*. in *S. cerevisiae*). These observations across all organisms, mirror and greatly expand recent observations made on bacterial and phage JDPs that also revealed the diversity of architectures among class C JDPs, more often than not with yet unknown cellular functions^40^.

We then analyzed the predicted subcellular localization of JDPs to assess if the classes are differently represented in various cellular compartments. Analysis of the organellar makeup of the dataset revealed that mitochondrial JDPs were mainly of class A and C (Supplementary Fig.2B) while mitochondrial class B JDPs were only about 10% of the class A JDPs, compatible with the precision of the localization algorithm itself (see Methods). This suggested that mitochondrial JDPs largely belong to class A, with some exceptions that must be manually inspected. ER-localized JDPs showed an intriguing class distribution. In addition to several class C JDPs, the ER contains a resident class A/B-type JDP, which in fungi (and some other unicellular eukaryotes such as amoebas) is a canonical class A, while in metazoa and plants (DNAJB11/Erdj3) it belongs to class B due to the lack of a ZFLR. A closer inspection revealed that, instead of the customary ZFLR, a non-canonical cysteine-rich region (which we call here *pseudoZFLR*) is generally found in these class B-like members. The pseudoZFLR regions display regularity and high degree of similarity in plants and metazoa (Fig.2G), with only two Cys pairs. The first Cys pair shows a highly conserved CNC motif, while the 2^nd^ pair has the two cysteines separated by two residues. This motif represents an interesting evolutionary conundrum, given that the most accepted view of the evolution of eukaryotes posits that the divergence in opisthokonts, between fungi and metazoan, took place after the split from plants.

### J-domain sequences contain functional and phylogenetic information at multiple scales

Organisms ranging from viruses to prokaryotes to eukaryotes rely on multiple pairs of Hsp70s and JDPs to support various biological activities. What largely remains unclear is how a particular JDP recognizes its partner Hsp70, an interaction that is largely mediated by the J-domain. It has been speculated for many years that the specificity should thus be encoded in the sequences of both partners, calling into question the degree to which the J-domains can be interchanged between different JDPs with minimal consequences for its primary function. In fact, not all J-domains interact indifferently with Hsp70s. Exchanges of J-domains between two JDPs frequently result in non-functional chimeras, suggesting that there is selectivity in the pairing of JDPs and Hsp70s^28,41–45^. Evolutionary and functional factors such as phylogeny, interactions with different Hsp70 paralogs, as well as the colocalization of J- and non-J-domains in the same polypeptide, may have left traces in the sequence of the J-domain. If this was the case, the sequence of a J-domain alone should carry relevant information about the overall JDP to which it belongs.

To test this hypothesis and provide evidence to support that the sequences of J-domains carry a rich source of disparate evolutionary and functional signatures, we first extracted and aligned 223’124 J-domain sequences from our JDP repertoire (see Methods for detailed procedure). We then used the unsupervised UMAP algorithm^46^ to provide a (non-linear) projection of the sequences in two dimensions, allowing for their direct visual inspection (Fig.3; see Methods). The projections revealed that the J-domain sequences are gathered in clusters, and that the different groups, highlighted by different colors, often associated well with various classification schemes, such as their *i*) A, B, or C class (Fig.3A), *ii*) bacteria rather than eukaryotes (Fig.3B) *iii*) various C subclasses (only the five more abundant ones are represented in Fig.3C for sake of clarity). To highlight the finer signatures that are encoded in the J-domain sequences, we focus on HscB. Besides being well-clustered according to the previous criteria (circles in Figs.3A, B, and C), the HscB sequences from fungi, metazoa and plants were also clearly individually well-grouped (Fig.3D). Thus, different and non-dependent layers of information are simultaneously encoded in the sequences of the J-domains, both to finer phylogenetic levels or for other criteria, such as subcellular localization (Supplementary Fig.4).

**Figure 3.**
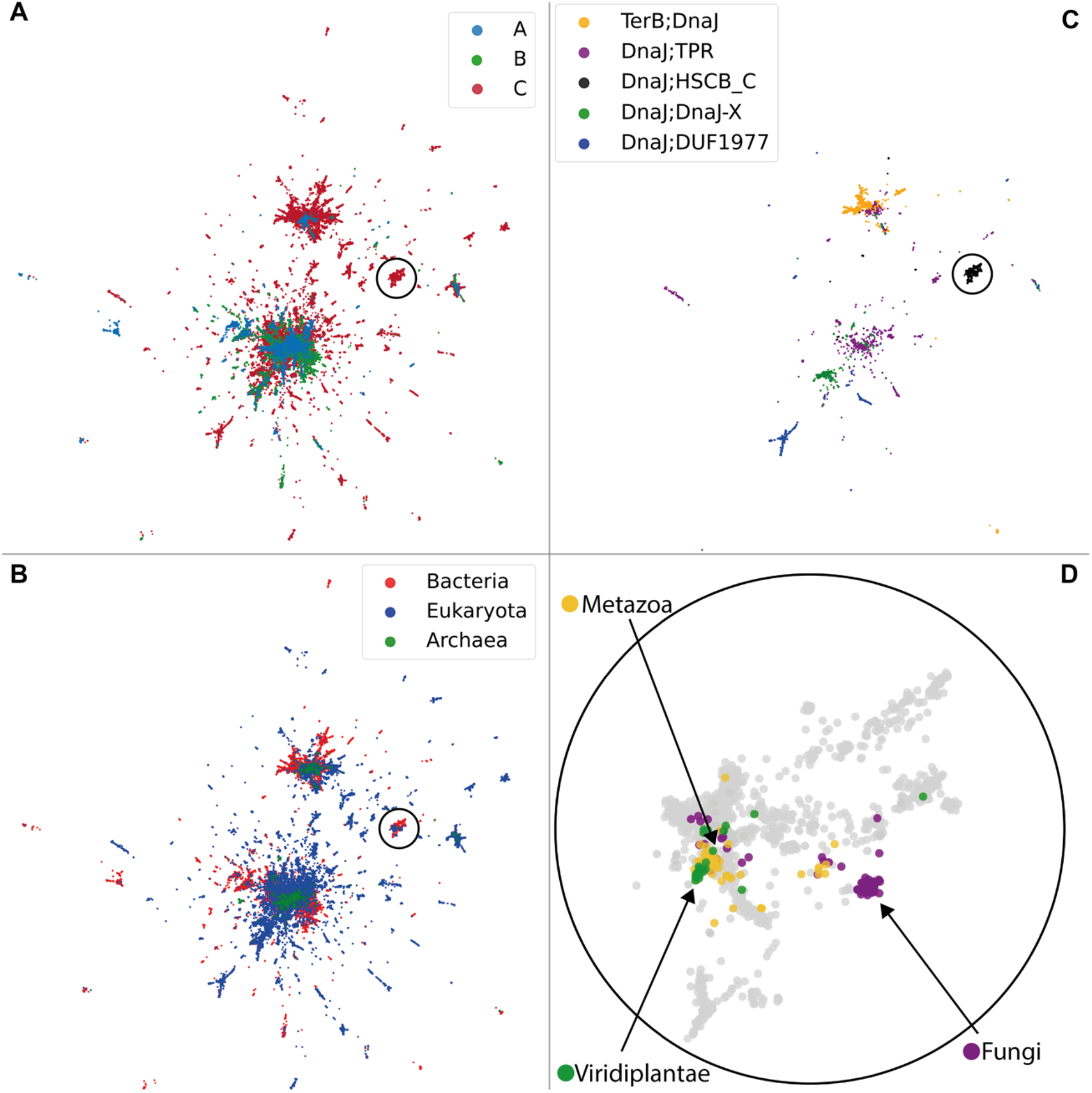
Two-dimensional UMAP representations of J-domains. JDP sequences are projected in two dimensions by a non-linear transformation (UMAP) and colored according to different schemes. **A)** J-domains are identified as being of class A (green), B (blue) or C (red) JDPs. **B)** J-domains are identified as being of bacterial (red) or eukaryotic (blue) origin. **C)** The five most abundant C subclasses are highlighted (see Fig.2F for representative members). The circles in each panel focus on the group identified as HscB-like JDPs. **D)** Zoom on the encircled region in the 3 other panels, highlighting the further differentiation of the eukaryotic ones into fungi, metazoa and plants (bacterial HscB sequences are shown in grey background). For visual clarity, a random subset of 50% of all points are displayed.

### Classification of J-domains by artificial neural networks

Next, we used an approach based on artificial neural networks (ANN) to find the detailed determinants in J-domain sequences that underpin different classifications. We focus on a few broad category discriminations, namely:

- Class A *vs*. class B (henceforth referred to as A/B);
- A *vs*. B *vs*. C (henceforth referred to as A/B/C);
- 12 different architectural classes, including the architectures of classes A and B JDP, the 9 most populated C subclasses (excluding JDPs containing only the DnaJ domain) and an additional “Other” class generically comprising all remaining JDPs (again not including JDPs containing only DnaJ domain) (henceforth referred to as Arch12). The 12 architectural classes in the Top12 task are thus “ClassA (DnaJ-DnaJ_C-DnaJ_CXXCXGXG”, “Class B (DnaJ-DnaJ_C)”, “TerB_DnaJ”, “DnaJ-TPR”, “DnaJ-HSCB_C”, “DnaJ-DUF1977”, “DnaJ-DnaJ-X”, “DnaJ-DnaJ_like_C11_C”, “DnaJ-zf-CSL”, “DnaJzf-C2H2_jaz”, “DnaJ_Sec63”, “Other”;
- Bacteria *vs*. eukaryotes *vs*. archaea *vs*. Viruses (henceforth referred to as Bac/Euk/Arch/Vir);
- Alphaproteobacteria *vs*. gammaproteobacteria *vs*. firmicutes *vs*. metazoa *vs*. plants *vs*. fungi (henceforth referred to as Phylo);
- Endoplasmic reticulum *vs*. mitochondria *vs*. plastids (henceforth referred to as Localization).

Although these different groupings are far from exhaustive, we chose them as representatives of both complementary and orthogonal criteria.

ANNs are often considered black-boxes with high performance for their assigned tasks, but poor interpretability. To circumvent this limitation, we employed a mathematical approach that makes ANNs interpretable, thus gaining a deeper insight into the information encoded in J-domain sequences. The sequence of operations performed by ANNs is shown in Fig.4A (and further detailed in the Methods).

**Figure 4.**
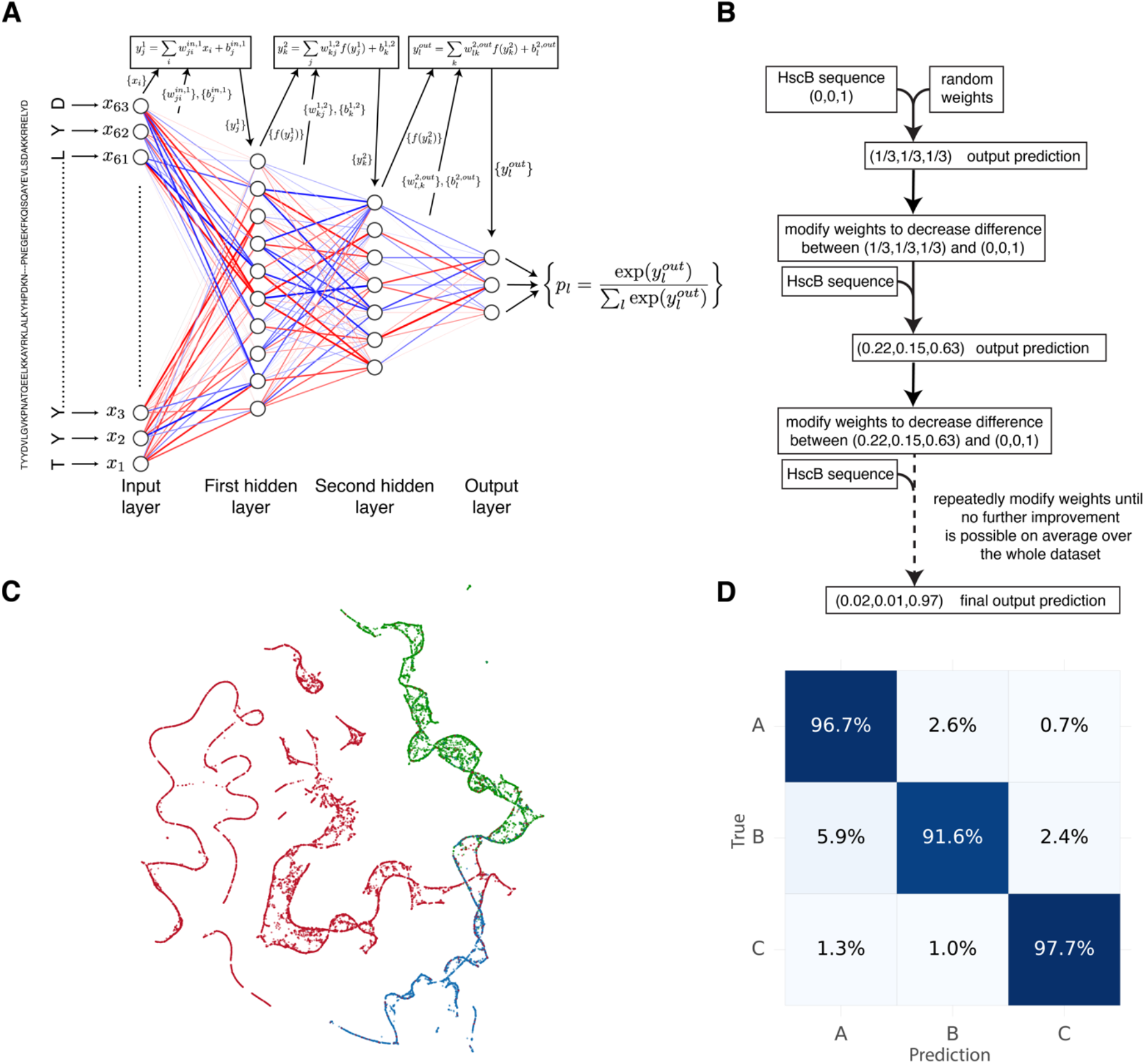
Structure, training and interpretation of Artificial Neural Networks. **A)** Structure of an ANN composed by input and output layers, separated by hidden layers (in this examples, two hidden layers); each node of a layer receives as input linear combinations of the values contained in the nodes of the previous layer, applies to them a non-linear function and the procedure is then repeated toward the next layer; the output layer transforms the received values into the probabilities for the input to belong to the various classes that must be distinguished. For visual clarity, the one-hot-encoding of the input sequences is not shown. The actual input size of all networks is 63*21=1323. **B)** Outline of the iterative procedure to change the ANN weights so that, on average over the training set, sequences are assigned as correctly as possible to their ground-truth class (defined according to domain composition). **C)** Two-dimensional embedding of the learnt probabilities for each J-domain sequence for the A *vs*. B *vs*. C classification task (colors according to the ground truth), providing a pictorial representation of the quality of the distinction of the classes. **D)** Confusion matrix for the A *vs*. B *vs*. C classification task: each entry tells what fraction of a given class is classified to the possible classes, providing an intuitive appraisal of the ANN performance.

The nodes (neurons) of the input layer receive the J-domain sequences in a suitable mathematical encoding, {*x*_*i*_}. The nodes of the first *hidden* layer receive *affine transformations* 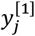 of the input values, with weights 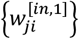 and offsets 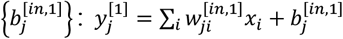 (subscripts indicate the connected nodes in the two layers, and the superscripts in the square brackets indicate corresponding layers). The output of the first hidden layer is then computed by applying a non-linear function to each of the 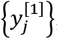, resulting in a set of values 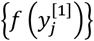, which is then passed to the nodes of the second hidden layer after undergoing the same set of operations. This procedure is repeated from layer to layer until the transferred values reach the nodes of the output layer that are in numbers equal to the different classes to be recognized (*e*.*g*. 3 output nodes for A/B/C, or 13 output nodes for Arch12). The output values are eventually transformed into the predicted probability that the input sequence {*xi*} belongs to each given class.

The training of the ANNs consisted in progressively modifying the network parameters (weights and offsets) until the output probabilities of the sequences of the dataset were, on average, as similar as possible to ground truths established previously (Fig.4B). For example, in the A/B/C classification task, the J-domain of HscB belongs to class C with certainty, corresponding to the probabilities (0,0,1), which is the set of values that the ANN should reproduce. To prevent the networks from overfitting by learning each sequences labels, the dataset was split into two parts. One part was used for training, while the other (test set) was used to check the predictions on sequences not seen during the learning phase. An ANN could be reliably used as a classification tool only if the performance on the training set was comparable to that on the test set, which we observed.

For each of the tasks (A/B, A/B/C, *etc*.), the output probabilities were used as coordinates for a low-dimensional representation of the dataset, similarly to the analysis performed in Fig.3. As an example, this kind of representation is shown in Fig.4C for the classification task A/B/C (including a further reduction in dimensionality from three to two dimensions using the UMAP algorithm), providing evidence that the ANN had found information embedded in J-domain sequences that allowed it to easily distinguish the three classes.

To move beyond the visual (but not quantitative) appraisal of the performance of the ANNs, we computed their *confusion matrices* (Fig.4D), which report the misclassification of each category with respect to their *ground truth*. An example is shown in Fig.4D where J-domains belonging to JDPs of class A were correctly classified 96.7% of the times, and misclassified as class B or C members only 2.6% and 0.7% of the times, respectively. Similarly, class B J-domains were classified according to their ground truth 91.6% of the times. Overall, the performance of the ANNs for all classification tasks was excellent (Supplementary Fig.5). The only classification task that was slightly less accurate was localization, although it is important to recall that the accuracy in this case was compatible with the prediction accuracy of the algorithms used to define the localization ground truth (see Methods). Overall, the success of our ANN classification approach highlights that a myriad of phylogenetic and functional signatures is indeed encoded in J-domain sequences alone and that this methodology can be used to tell J-domains apart from each other depending on different, possibly orthogonal, criteria such as phylogeny, subcellular localization, and domain organization. The co-evolution of J-domain sequences with the full protein architecture has also been recently reported, albeit restricted to bacterial genomes^40^.

We finally used our trained ANNs to predict the A, B, and C classes, as well as the Archs12 labels of the 78’282 JDPs with an annotated domain architecture composed of a single DnaJ. Note again that none of these ∼78’000 sequences were used during network training. The ANNs correctly classified vast majority of these as class C JDPs. Interestingly, more than 60% of these singleton JDPs were classified as “Other” in the Archs12 task, implying that the network has probably learned that the divergent signatures of these JDPs are dissimilar from the other classes used to learn this task. The second and third most predicted classes on this set correspond to class A and B JDPs (∼9% and ∼17% respectively), which is likely due to them being the most abundant JDPs in the training set. Collectively, these results seem to indicate that a substantial fraction of JDPs with the J-domain alone, or together with unannotated co-domains, exist, and that these form a (probably heterogenous) divergent clade of J-domains.

### Structural characterization of the most relevant discriminative sequence positions

Next, we used the interpretability of the ANNs to identify the sequence positions that were most relevant for each classification task (see Methods) by assigning them a *relevance score*^47^. We found that although several of the sequence positions were expectedly task-specific, some were ubiquitously important for all tasks (heatmap in Fig.5A). In general, the average score of the relevance for different tasks highlighted the positions that were most relevant to classify J-domains. Thus, we map the five top scoring positions in the structure of the J-domain of *E. coli* DnaJ in the DnaJ-DnaK chaperone complex (*E. coli* Hsp70) chaperone complex (Fig.5B; the five most relevant positions for each classification task are shown in Supplementary Fig.6)^21^. Importantly, we observed that the five top-scoring positions, located at helix II and III of the J-domain, overlapped with the interaction surface between the J-domain of DnaJ and DnaK^21^. This is an intriguing result, because at no step of our analysis we used information about the binding of J-domains to Hsp70. It is thus tempting to hypothesize that the various, possibly orthogonal, classifications are successful because, among other factors, J-domains coevolved together with Hsp70 in a concerted phylogenetic and functional divergence. We presume that these positions likely facilitate modulation of the interaction between J-domain and Hsp70, and HPD-dependent tuning of Hsp70’s ATPase activity.

**Figure 5.**
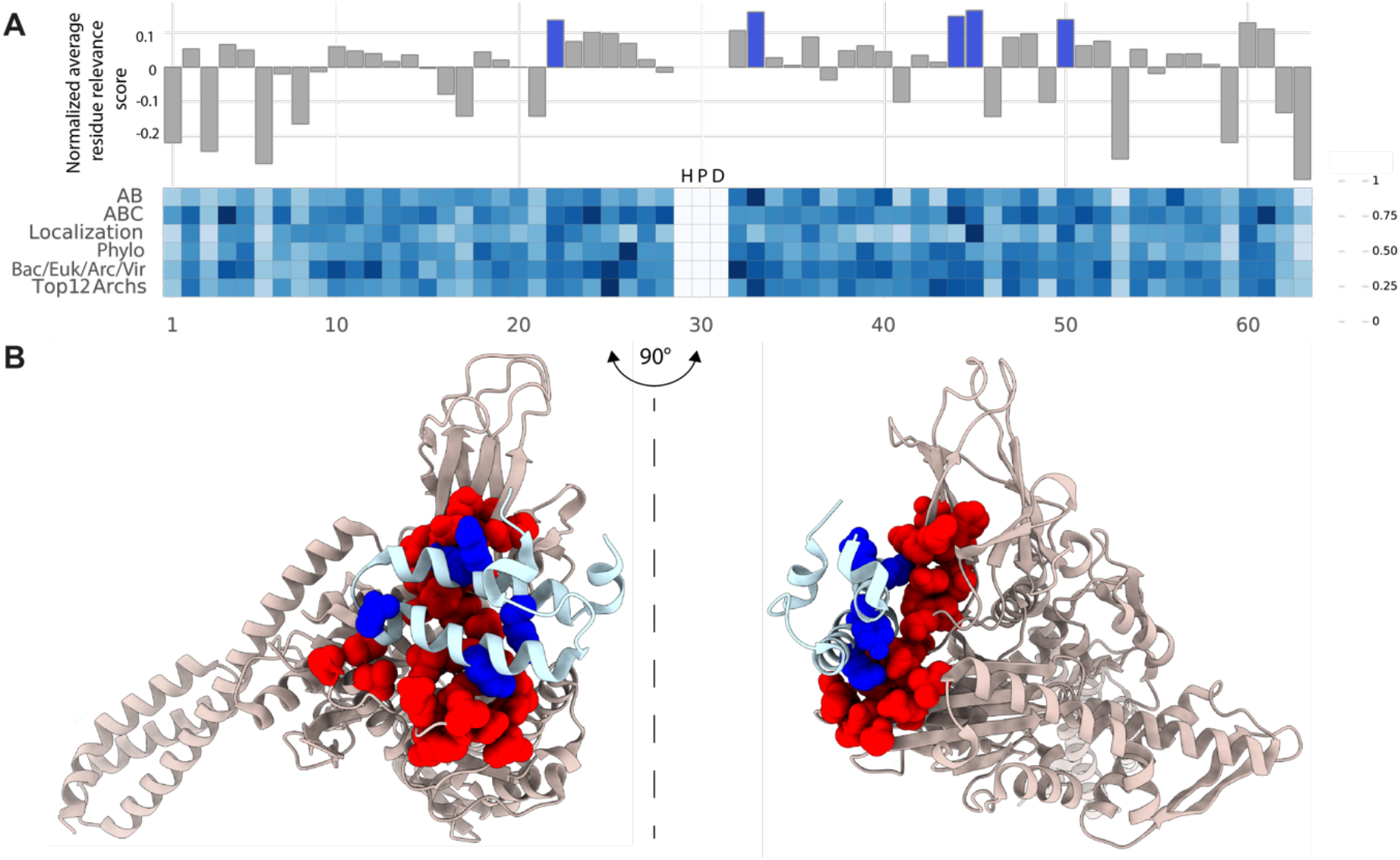
Identification of the most sequence and structure positions relevant for the classification. **A)** The relevance score for each position and for different classification tasks is represented in a heatmap. For each position, the relevance score averaged over the different tasks is represented, relative to the average over all positions, above the heatmap (HPD are not considered as they are completely irrelevant for all tasks). The top five positions are highlighted in blue. **B)** The five top scores identified in the top panel are highlighted in blue on the structure of the J-domain/DnaK complex^21^, while the corresponding contacting residues on DnaK are represented in red, showing that the five most relevant positions on the J-domain participate in the interaction surface between the two proteins.

### Classification of J-domain Proteins according to their J-domains

Next, we employed the ANNs as class predictors for the human JDPs. We first excluded all human JDPs, together with those of other model organisms (see Methods) from the training set to prevent any bias during the learning process. In Table 1, we report the probabilities attributed to each human J-domain with respect to a given class or architectural group.

**Table 1.** Classification of the 50 *H. sapiens* JDPs. Architectures grouped into the “Other” class are indicated by *(Other)* in the third column. Class C JDPs mis-predicted in the Arch12 task are highlighted in orange: DNAJC20 is actually recognized as a HscB class C protein, although the presence of further domains in the human homolog makes the classification formally wrong; a similar argument applies to DNAJCC23 (SEC63). The presence of corrupted HPDs (*e*.*g*. DNAJB13) does not change the classification based on the J-domain, because all J-domains used in the training of the Arch12 ANN had a canonical HPD motif. As a consequence, those positions are immaterial to the classification. For non-canonical class B JDPs, we report in green the modified nomenclature introduced in^28^. The reference proteome (UP000005640 (Homo sapiens)) was used to perform the search of the JDPs.

|  | *H. Sapiens* name | Uniprot | Domain Architecture (Ground truth) | A/B/C Probabilities | Arch12 Probabilities |
| --- | --- | --- | --- | --- | --- |
| **Class A** |  |  |  |  |  |
| 1 | DNAJA1 | P31689 | DnaJ - DnaJ_C - DnaJ_CXXCXGXG | **0.94** / 0.06 / 0.00 | DnaJ;DnaJ_C;DnaJ_CXXCXGXG (**0.98**) |
| 2 | DNAJA2 | O60884 | DnaJ - DnaJ_C - DnaJ_CXXCXGXG | **1.00** / 0.00 / 0.00 | DnaJ;DnaJ_C;DnaJ_CXXCXGXG (**1.0**) |
| 3 | DNAJA3 | Q96EY1 | DnaJ - DnaJ_C - DnaJ_CXXCXGXG | **0.99** / 0.00 / 0.00 | DnaJ;DnaJ_C;DnaJ_CXXCXGXG (**0.97**) |
| 4 | DNAJA4 | Q8WW22 | DnaJ - DnaJ_C - DnaJ_CXXCXGXG | **0.89** / 0.10 / 0.01 | DnaJ;DnaJ_C;DnaJ_CXXCXGXG (**0.96**) |
| **Canonical Class B** |  |  |  |  |  |
| 5 | DNAJB1 | P25685 | DnaJ - DnaJ_C | 0.00 / **1.00** / 0.00 | DnaJ;DnaJ_C (**0.98**) |
| 6 | DNAJB4 | Q9UDY4 | DnaJ - DnaJ_C | 0.00 / **1.00** / 0.00 | DnaJ;DnaJ_C (**1.0**) |
| 7 | DNAJB5 | O75953 | DnaJ - DnaJ_C | 0.00 / **0.99** / 0.01 | DnaJ;DnaJ_C (**0.98**) |
| 8 | DNAJB11 | Q9UBS4 | DnaJ - DnaJ_C - pseudoZFLR (*Other*) | 0.00 / 0.04 / **0.96** | Other (**0.89**) |
| 9 | DNAJB13 (not HPD) | P59910 | DnaJ - DnaJ_C | 0.00 / 0.01 / **0.99** | DnaJ;DnaJ_C (**0.77**) |
| **Non-canonical Class B** |  |  |  |  |  |
| 10 | DNAJB2 (DNAJC31) | P25686 | DnaJ | 0.00 / 0.03 / **0.96** | DnaJ;DnaJ_C (**0.92**) |
| 11 | DNAJB3 (DNAJC32) | Q8WWF6 | DnaJ | **0.45** / 0.11 / 0.44 | DnaJ;DnaJ_C (**0.62**) |
| 12 | DNAJB6 (DNAJC33) | O75190 | DnaJ | **0.58** / 0.05 / 0.37 | DnaJ;DnaJ_C;DnaJ_CXXCXGXG (**0.63**) |
| 13 | DNAJB7 (DNAJC34) | Q7Z6W7 | DnaJ | **0.76** / 0.11 / 0.14 | DnaJ;DnaJ_C;DnaJ_CXXCXGXG (**0.79**) |
| 14 | DNAJB8 (DNAJC35) | Q8NHS0 | DnaJ | 0.03 / 0.01 / **0.97** | DnaJ;DnaJ_C;DnaJ_CXXCXGXG (**0.58**) |
| 15 | DNAJB9 (DNAJC36) | Q9UBS3 | DnaJ | 0.12 / 0.00 / **0.88** | DnaJ;DnaJ_C;DnaJ_CXXCXGXG (**0.82**) |
| 16 | DNAJB12 (DNAJC37) | Q9NXW2 | DnaJ - DUF1977 | 0.03 / 0.00 / **0.96** | DnaJ;DUF1977 (**1.0**) |
| 17 | DNAJB14 (DNAJC38) | Q8TBM8 | DnaJ - DUF1977 | 0.00 / 0.00 / **1.00** | DnaJ;DUF1977 (**1.0**) |
| **Class C** |  |  |  |  |  |
| 18 | DNAJC1 | Q96KC8 | DnaJ - Myb_DNA-binding (*Other*) | 0.00 / 0.00 / **1.00** | Other (**1.0**) |
| 19 | DNAJC2 | Q99543 | DnaJ - RAC_head - Myb_DNA-binding - Myb_DNA-binding (*Other*) | 0.00 / 0.00 / **1.00** | Other (**1.0**) |
| 20 | DNAJC3 | Q13217 | TPR_16 - TPR1 - TPR_8 - DnaJ | 0.00 / 0.00 / **1.00** | DnaJ;TPR (**1.0**) |
| 21 | DNAJC4 | Q9NNZ3 | DnaJ | 0.00 / 0.00 / **1.00** | Other (**1.0**) |
| 22 | DNAJC5 | Q9H3Z4 | DnaJ | 0.00 / 0.01 / **0.99** | Other (**0.97**) |
| 23 | DNAJC5B | Q9UF47 | DnaJ | 0.00 / 0.00 / **1.00** | Other (**0.63**) |
| 24 | DNAJC5G | B5MCA9 | DnaJ | 0.00 / 0.00 / **1.00** | Other (**0.98**) |
| 25 | DNAJC6 (AUXILIN) | O75061 | PTEN_C2 - DnaJ (*Other*) | 0.00 / 0.00 / **1.00** | Other (**1.0**) |
| 26 | DNAJC7 | Q99615 | TPR_11 - TPR_16 - TPR_8 - DnaJ | 0.02 / 0.03 / **0.95** | DnaJ;TPR (**1.0**) |
| 27 | DNAJC8 | O75937 | DnaJ | 0.00 / 0.00 / **1.00** | Other (**1.0**) |
| 28 | DNAJC9 | Q8WXX5 | DnaJ | 0.00 / 0.00 / **1.00** | Other (**1.0**) |
| 29 | DNAJC10 | Q8IXB1 | DnaJ - Thioredoxin - Thioredoxin - Thioredoxin - Thioredoxin (*Other*) | 0.00 / 0.00 / **1.00** | Other (**0.99**) |
| 30 | DNAJC11 | Q9NVH1 | DnaJ - DnaJ-like_C11_C | 0.00 / 0.00 / **1.00** | DnaJ;DnaJ-like_C11_C (**1.0**) |
| 31 | DNAJC12 | Q9UKB3 | DnaJ | 0.00 / 0.00 / **1.00** | Other (**1.0**) |
| 32 | DNAJC13 | O75165 | RME-8_M - GYF_2 - DnaJ (*Other*) | 0.00 / 0.00 / **1.00** | Other (**1.0**) |
| 33 | DNAJC14 | Q6Y2X3 | DnaJ - Jiv90 (*Other*) | 0.00 / 0.00 / **1.00** | Other (**1.0**) |
| 34 | DNAJC15 | Q9Y5T4 | DnaJ | 0.00 / 0.00 / **1.00** | Other (**1.0**) |
| 35 | DNAJC16 | Q9Y2G8 | DnaJ - Thioredoxin | 0.05 / 0.03 / **0.92** | Other (**0.61**) |
| 36 | DNAJC17 | Q9NVM6 | DnaJ - RRM_1 (*Other*) | 0.00 / 0.00 / **1.00** | Other (**1.0**) |
| 37 | DNAJC18 | Q9H819 | DnaJ - DUF1977 | 0.00 / 0.00 / **1.00** | DnaJ;DUF1977 (**1.0**) |
| 38 | DNAJC19 (TIM14) | Q96DA6 | DnaJ | 0.00 / 0.00 / **1.00** | Other (**1.0**) |
| 39 | DNAJC20 (HSCB) | Q8IWL3 | HscB_4_cys - DnaJ - HSCB_C | 0.00 / 0.00 / **1.00** | DnaJ;HSCB_C (**0.99**) |
| 40 | DNAJC21 | Q5F1R6 | DnaJ - zf-C2H2_jaz | 0.00 / 0.00 / **1.00** | DnaJ;zf-C2H2_jaz (**1.0**) |
| 41 | DNAJC22 | Q8N4W6 | TM2 - DnaJ (*Other*) | 0.00 / 0.00 / **1.00** | Other (**1.0**) |
| 42 | DNAJC23 (SEC63) | Q9UGP8 | DnaJ - Sec63 - Sec63 (*Other*) | 0.00 / 0.00 / **1.00** | DnaJ;Sec63 (**0.97**) |
| 43 | DNAJC24 | Q6P3W2 | DnaJ - zf-CSL | 0.00 / 0.00 / **1.00** | DnaJ;zf-CSL (**1.0**) |
| 44 | DNAJC25 | Q9H1X3 | DnaJ | 0.00 / 0.00 / **1.00** | Other (**0.95**) |
| 45 | DNAJC25-GNG10 | A0A024R161 | DnaJ - G-gamma (*Other*) | 0.00 / 0.00 / **1.00** | Other (**0.98**) |
| 46 | DNAJC26 (GAK) | O14976 | Pkinase - PTEN_C2 – DnaJ (*Other*) | 0.00 / 0.00 / **1.00** | Other (**1.0**) |
| 47 | DNAJC27 | Q9NZQ0 | Ras – DnaJ (*Other*) | 0.00 / 0.00 / **1.00** | Other (**1.0**) |
| 48 | DNAJC28 | Q9NX36 | DnaJ – DJC28_CD (*Other*) | 0.00 / 0.00 / **1.00** | Other (**1.0**) |
| 49 | DNAJC29 (SACSIN) | Q9NZJ4 | Ubiquitin – DnaJ – HEPN (*Other*) | 0.00 / 0.00 / **1.00** | Other (**0.98**) |
| 50 | DNAJC30 | Q96LL9 | DnaJ | 0.00 / 0.00 / **1.00** | DnaJ;DnaJ-X (**1.0**) |

The J-domains of human class A (DNAJA1-4) and “canonical” class B (DNAJB1, DNAJB4, DNAJB5) were correctly classified according to the tasks A/B, A/B/C and Arch12 (Table 1 and Supplementary Table 2 for the task A/B), confirming that the ANNs were able to reliably learn their characteristics during the training phase. A more specific discussion is warranted for DNAJB11 and DNAJB13, whose J-domains were consistently classified as class C according to the A/B/C classification task. They were, however, correctly classified by the Arch12 task as either “Other” (DNAJB11) or recognized as canonical class B in the Arch12 task (Table 1). These results suggest that DNAJB11 and DNAJB13, despite a likely class B origin, have further diverged enough that their sequences do not fully conform to the majority of class B JDPs.

**Table 2.**
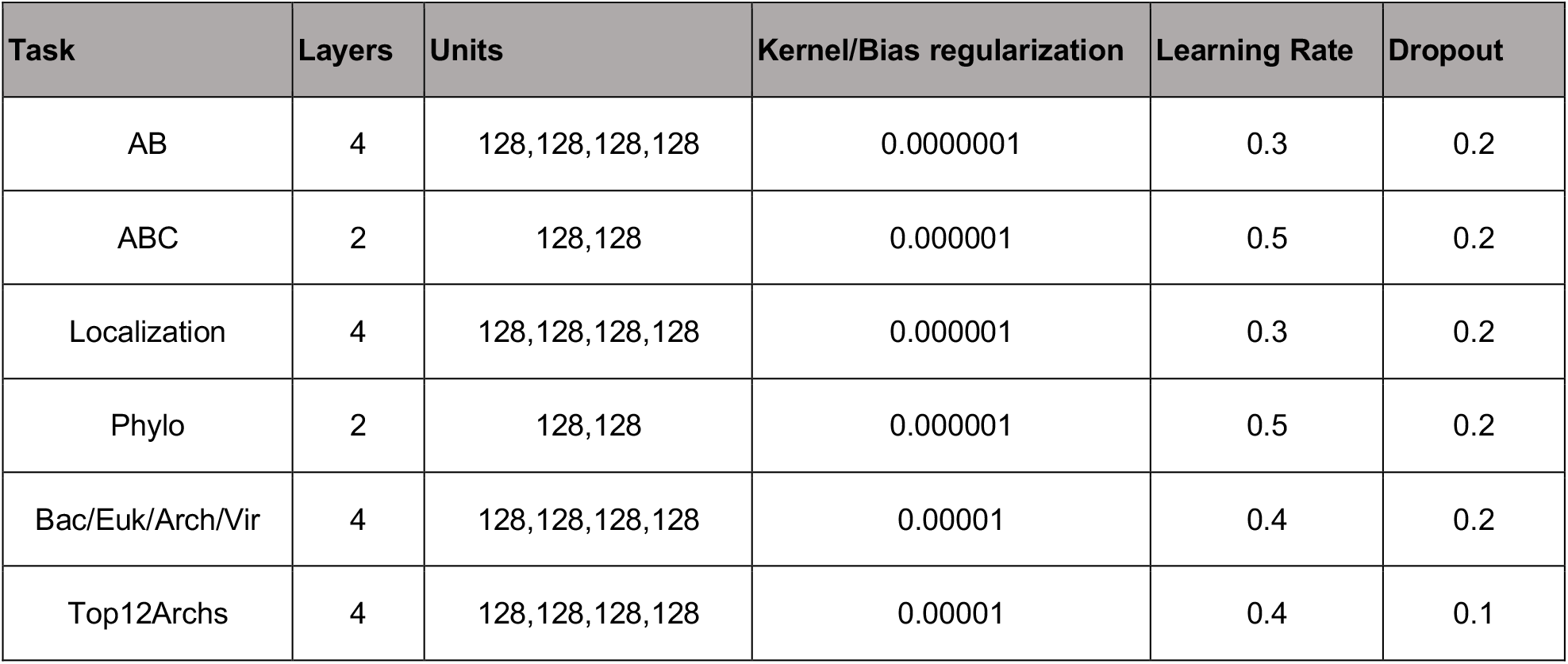
Optimal hyperparameters used for training the different classification networks.

Class B JDPs with non-canonical architectures (DNAJB2, DNAJB3, DNAJB6, DNAJB7, DNAJB8, DNAJB9, DNAJB12, and DNAJB14) have historically been attributed to class B due to the presence of an N-terminal J-domain followed by a G/F-rich region. We used both the A/B/C and Arch12 classification tasks to try and reclassify these proteins. According to the A/B/C task, none of them should be considered genuine class Bs. We then cross-checked this prediction with the Arch12 task. We found that DNAJB12 and DNAJB14 belonged to class C and in particular to the same architectural Class as DNAJC18, with whom they share the DUF1977 domain, while the other non-canonical class B JDPs were predicted to belong to either class A or class B. A smaller model discriminating solely class A from class B JDPs yielded consistent results (see Supplementary Table 2). Taken together, these results suggest a complex evolutionary origin of the non-canonical class B JDPs, with some of them possibly descending from class A JDPs (Table 1).

Next, we classified class C JDPs into subclasses. All original class C members were correctly predicted by the A/B/C classification task. The Arch12 task also correctly classified all human class C JDPs. Of note, DNAJC20 (HscB) and DNAJC23 (Sec63) (highlighted in orange in Table 1) were assigned to the *Others* group, since these JDPs displayed domain architectures somewhat divergent from the Pfam-annotated domains. However, the Arch12 task correctly assigned DNAJC20 and DNAJC23 to HscB and Sec63 subclasses, highlighting the ability of these automated approaches to correctly annotate JDPs with the help of appropriate training sets. We repeated this analysis to several other model organisms covering a wide phylogenetic range, including *M. musculus, R. norvegicus, M. mulata, D. rerio, D. melanogaster, C. elegans, A. thaliana* and *E. coli* (see Supplementary tables S3-S10).

## Discussion

In the course of evolution, the JDP family expanded both in number and structural diversity, and in eukaryotic organisms there is a more than two-fold increase in the number of JDPs relative to Hsp70s, indicating that the limited repertoire of Hsp70 isoforms is utilized by multiple different JDPs to generate various chaperoning activities (*e*.*g*. unfolding of misfolded protein, disassembly of functional or aggregated protein oligomers, protein translocation into organelles and target proteins for degradation). The assembly of different JDP-Hsp70 chaperone machines is initiated by J-domains. Genetic, biochemical, and structural data accumulated over the years provide a relatively comprehensive map of the J-domain-Hsp70 interface^20,21,48–53^, which serves a single task, *i*.*e*. to establish the vital physical interaction that promotes ATP hydrolysis in Hsp70^16,17,20–23^, thus allowing the chaperone to capture client substrates^18^. Proper docking of the J-domain onto ATP-bound Hsp70 is ensured by a core set of intermolecular contacts occurring along the helix II--loop (HDP motif)--helix III region of the J-domain, which is structurally conserved from bacteria to human^20,21^. However, there is very little knowledge of the regulatory elements in the J-domains that critically fine-tune this interaction^41^ despite such modulation being hypothesized over the years. Our ANN-based observations show the presence of phylogenetic and functional signatures buried within the amino acid sequences of J-domains that appear to modulate the pairing of JDPs with Hsp70s to form chaperone machines with different activities.

What is the basis for this functional and phylogenetic sorting? Remarkably, the most discriminative sequence positions structurally coincided with the J-domain-Hsp70 binding interface, strongly hinting that some amino acids have a regulatory role to optimize the selection of unique functionally specific Hsp70-JDP pairs. These *in silico* observations are strongly supported by indirect experimental evidence. For example, measurements of JDP stimulated ATPase activity in Hsp70s shows that J-domains are not fully interchangeable with respect to their collaboration with different Hsp70s within a given species. The yeast cytosol contains seven Hsp70 paralogs (Ssa1-4, Ssb1-2 & Ssz1). Both Ssa’s and Ssb’s bind to nascent polypeptide chains to promote *de novo* protein folding. However, only the ATPase activity of Ssa’s, but not of Ssb’s, is stimulated *in vitro* by the two major cytosolic yeast JDPs Ydj1 (class A) and Sis1 (class B)^54^. This indicates that J-domains of Ydj1 and Sis1 have evolved to differentially recognize these Hsp70 family members. Similarly, the ATPase activity of ER-localized yeast Hsp70 Bip/Kar2 (HSPA5) was shown to be induced by the J-domain of Sec63 (ER; class C) 16 times higher than the J-domain of ERdj3 (ER; class C)^55^ suggesting that, though both JDPs function with this Hsp70, there are differences in how the two JDPs stimulate Bip for specific activities. Recent results have also highlighted that, for example, the J-domain of human DNAJC27 involved in ciliogenesis, cannot be substituted by almost any other human J-domain^56^. In addition, as expected, speciation also has left a clearly discernible signature on the J-domain sequence, which points towards some incompatibilities in JDP-Hsp70 pairing. Pairing of Hsp70s and JDPs from different organisms have generated poor chaperoning activities. For example, bacterial class A JDP DnaJ stimulates the ATPase activity of bacterial DnaK and human HSPA8 (Hsc70) very differently^57^. Similarly, mouse DNAJC7 (DjC7; class C JDP) stimulates the ATPase activity of bacterial DnaK and murine Hsp70s (HSPA8 and HSPA5) at varying levels^58^ possibly leading to differences in function. Our *in silico* findings strongly suggest that these incompatibilities arise from discriminatory signatures buried within the amino acid sequences of J-domains. Emergence of greater sequence diversity in J-domains of JDPs, in particular at the Hsp70-J-domain interaction surface, with respect to their Hsp70 partners is expected to preserve specificity in pairing as the Hsp70-JDP network expanded during evolution. In general, amino acid changes at the relatively small Hsp70-J-domain interaction surface would require independent compensatory mutations in both the J-domain and Hsp70 to maintain compatibility. However, this is statistically very unlikely, because it imposes a greater evolutionary pressure on the handful of Hsp70s (compared to the large number of JDPs) in a cell. On the contrary, there is relatively less selection pressure for mutations to occur on the J-domain side, but with a certain affinity cost for the pairing. As a result, we observe the establishment of a gradient of interaction preferences between distinct Hsp70 and JDP pairs. This largely explains the aforementioned J-domain involved Hsp70-JDP incompatibilities reported experimentally. Overall, the evolution of these discriminatory genetic signatures through mutations in J-domains opens the possibility to fine-tune JDP-Hsp70 interactions to support specific chaperoning activities in cells.

Our analysis refined the JDP classification. The ANNs used show that the unique amino acid sequence signatures in the J-domains also provide precise discrimination according to functional and phylogenetic criteria. For example, the discriminatory signatures in the J-domains of non-canonical class B JDPs (DNAJB2, DNAJB3, DNAJB6-9, DNAJB12 and DNAJB14) suggest that these JDPs are very distinct from the canonical class B members such as DNAJB1, DNAJB4 and DNAJB5. This is further supported by large variations in the domain architecture of the full-length proteins. The presence of a G/F-rich region following the J-domain suggests that these JDPs may have evolved from a class A or class B ancestor, followed by some extensive sequence divergence. Class C appears to be the largest subfamily in prokaryotes^40^ and has exponentially increased in number during eukaryotic evolution, being further organized in subclasses that are clearly recognizable according both to their overall domain architecture and to the sequences of their J-domains alone.

From a more general perspective, the recent advent of ANN-based approaches in structural biology, epitomized by the spectacular and widely-advertised success of AlphaFold for protein structure prediction^59,60^, highlights that the exponentially growing number of sequences available in public repositories holds a yet unexplored wealth of information that must be exploited. At the same time, this data abundance also creates a problem between indiscriminate use of sequences from all sources and cautious desire to use carefully curated sequences, and calls for an intermediate approach, where stringent, conservative automated criteria go hand-in-hand with manual curation at higher scales (*e*.*g*. domain composition). In this respect, the present work can be considered as an important example where this strategy allows for an unbiased, semi-unsupervised coupled functional/phylogenetic-structural investigation: It is supervised for the classification task, but unsupervised for the emergence of the relevant structural features.

### Limitations of the study

We show here that the proposed automated classification is already possible, although likely limited mostly by two factors. On the one hand, several architectural C subclasses are too poorly represented in the dataset that it is difficult to train a network to tell them apart from more abundant subclasses. However, this problem is likely to be solved through exponential growth of the number of reported JDP sequences. On the other hand, a large swath of class C JDPs comprise domains that have not yet been annotated in Pfam, or just some disordered regions of various lengths. Yet, the advent of novel deep learning approaches on scales even larger than the present ones is recognizing domains that were previously uncharacterized^61^, paving the way for a better assignment of class C JDPs to precise, well-populated architectural classes.

## Methods

### Construction of a curated J-protein dataset

We constructed a JDP sequence database by querying the Uniprot database (union of Swissprot and TrEMBL, release 2021_04)^31^ to extract all sequences annotated to have a J-domain according to the PFAM domain annotation (PF00226)^32^. For each JDP sequence, we recovered its full-length sequence as well as the full taxonomic lineage from Uniprot. We then extracted the J-domain sequence of each JDP by aligning the full-length sequences to the Hidden Markov Model (HMM) of the Pfam J-domain (PF0226) family using the *hmmalign* utility of the *hmmer* package^62^. The full domain-architecture was obtained by querying the PFAM database for each JDP entry in our database and the domain order validated using the domain boundaries reported in PFAM.

The domain architectures were used to classify all sequences as either belonging to canonical class A (PFAM architecture “*DnaJ;DnaJ_C;DnaJ_CXXCXGXG*”), canonical class B (PFAM architecture “*DnaJ;DnaJ_C;*”) and class C (all other sequences).

We further identified the G/F-rich region in all JDPs as being the stretch of 40 amino-acids directly following the JD sequence, if at least 25% of its amino-acids are either Glycines or Phenylalanines. For ease of analysis, we also grouped all domains composing TPR repeat proteins (annotated as TPR_XX) into a virtual TPR domain. Thus, in all proteins containing one or several TPR repeats, we replaced all the TPR domains by a single effective TPR domain (as shown in Fig.2). Subcellular localizations were predicted for eukaryotic sequences using the TargetP software^63^, version 2.0.

This resulted in an initial dataset containing 232’516 annotated JDP sequences.

We then curated the dataset by first removing all sequences for which the Uniprot and PFAM annotations were not consistent and also removed all sequences which did not have a full characteristic HPD motif. We further filtered out sequences containing annotated ANTI-TRAP domains (according to Pfam), because in all but one case it simply recognized the canonical ZFLR domain. Furthermore, we also removed sequences containing multiple J-domains. Finally, all sequences associated to unclassified organisms were also removed from our final set.

This resulted in a curated JDP dataset comprising 223’124 sequences.

### Annotation of a novel pseudo Zinc-Finger Like Region (pseudoZFLR) domain in JDPs

We created a new domain definition to annotate the pseudo zinc-finger like domains similar to the ones observed in DnaJB11 and ErdJ3. These domains are characterized by one CNC and one CxxC motifs, in contrast to the 4 pairs of CxxC motifs of the canonical Zinc-Finger motif as annotated in the PFAM DnaJ_CXXCXGXG domain. We thus built a multiple sequence alignment containing 35 manually selected and curated JDPs containing a pseudoZFLR domain, from organisms including vertebrates, invertebrates and plants. We then aligned this seed using MAFFT (v7.487)^64^ and manually identified a region comprised of 55 positions defining the pseudoZFLR domain. We then constructed a hidden-markov model profile of the novel domain using the *hmmbuild* utility from the hmmer suite^62^ and used to profile to query all the 223’124 JDP sequences in our curated dataset to identify JDPs containing a pseudoZFLR domain. Because of the similarity between this domain and the canonical ZFLR domain, we used a more stringent E-value =10^−4^ cutoff to identify pseudo ZFLR domains (see Supplementary Fig.7 7). This resulted in a total of 1421 JDPs identified as containing a pseudoZFLR domain, of which 1336 possess a DnaJ-DnaJ_C-pseudoZFLR architecture.

### Dimensionality reduction of JDP amino-acid sequences by UMAP

To apply the UMAP algorithm^46^ to the amino-acid sequences of our J-domain repertoire for visualization, we first used a one-hot-encoding technique to numerically embed the JD sequences in a high-dimensional space, consisting of 63×21=1323 dimensions (20 amino-acid + 1 gap for each of the 63 positions in the JD aligned sequences). We then applied the UMAP dimensionality reduction technique with standard parameters as provided by the original publication.

### Preparation of JDP datasets for training Neural Networks

To avoid training the Neural Networks on sequences for which we want to predict and interpret the class labels, sequences from model organisms (*H. sapiens, M. musculus, S. cerevisiae, R. norvegicus, melanogaster, D. rerio, C. elegans, A. thaliana*) were removed from the dataset prior to training. To interpret the predictions of the trained models, we also removed non-canonical class B proteins from our training set. These were identified as JDPs containing a DnaJ domain, an annotated G/F region (see above) but no DnaJ_C domain. This procedure resulted in a reduced set of 205’252 sequences used for building the classification models.

To reduce the effects of sampling bias during the training phase, we used as simple sequence reweighting scheme previously used in several sequence analysis pipelines^65,66^. We assigned to each sequence a weight of 1/n_80_, where n_80_ denotes the number of sequences with sequence identity greater than 80% from it. These weights were calculated for each classification task separately, as some tasks are only trained on a subset of sequences, which can potentially slightly modify the weights.

We randomly split the processed dataset into training, validation and test set in such a way that sequences from the same organism always belong to the same set and that the sum of the weights of each dataset is respectively 0.7, 0.15, 0.15 times that of the total.

### Feed-Forward Artificial Neural Network classification

To feed the sequences to a neural network we encoded them into binary vectors. Every amino acid is mapped to an integer *A(a)* in increasing alphabetical order, with the gap symbol mapped to zero. We therefore can encode every amino acid with a 21-dimensional vector in which all elements are zeros and element *A(a)* is one (the so called *one-hot encoding*). An amino acid sequence is then obtained by concatenating all 21-dimensional vectors together. To pass the sequences to a neural network, we concatenated the one-hot encoding of each position in the alignment, resulting in a 1323-dimensional binary vector (63×21 entries).

A Dropout layer has been used between the input and first hidden layer to randomly mask a fraction of inputs and has been found to improve the generalization accuracy of the network.

For each task described in the main text, we trained a feed forward neural network to predict the corresponding label of each sequence. We trained the network by minimizing the categorical cross entropy between the output layer and the categorical encoding of the label. All weights have been regularized using an L2 regularizer. In all cases we used the Adam protocol during the minimization procedure and a learning decay equal to 0.01. A batch size of 1024 was used and SoftPlus activation were used for all hidden layers. The remaining hyperparameters (number of layers, units per layer, dropout rate, weight regularizations and initial learning rate) were optimized by performing a grid search over parameter space, testing 216 combinations of hyperparameters per task. The best performing model was selected based on the maximum accuracy on the validation set (see Table 2). The model performances are reported on the hold out test set.

After the training is completed, we analyzed the trained networks using the technique described in^47^ to obtain 21×63 coefficients describing how the networks use every bit in the input in order to classify the sequence. The relevance scores for each residue is finally computed as the Frobenius norm (sum of the squares) of its corresponding elements in the coupling matrix (see^47^ for details on the relevance score calculations).

### Class A vs B JDP classification using unaligned G/F region sequences

To test the discrimination power of the G/F regions of class A and B JDPs, we built an L_2_-regularized logistic regression model to classify JDPs as either belonging to class A or B. The unaligned G/F regions (comprised of the 40 amino-acids following the J-domain) where one-hot encoded and fed into a standard logistic regression model. The regularization parameter C has been chosen by scanning the validation set accuracy (Supplementary Fig.8). The resulting best model (C=3.0) achieved classification accuracy of ∼95%.

### Generate of J-protein repertoire for model organisms

We built the J-protein repertoires for the following model organisms: *H. sapiens, S. cerevisiae, M. musculus, R. norvegicus, M. mulata, D. rerio, D. melanogaster, C. elegans, A. thaliana* and *E. coli*. All entries in the human, yeast and bacterial organisms were manually curated and cross-referenced with J-protein repertoires of these organisms. For the remaining model organisms, we extracted all entries in our dataset correspondoing to the target organism, removed entries marked as fragment and then reduced redundancy by only retaining entries with both unique full-length sequences and gene names. To select which Uniprot entry to keep in the tables, we prioritized entries in the annotated SwissProt repository of Uniprot, and then selected the entries with the highest Uniprot annotation score.

## Author Contributions and Notes

DM and SZ conceptualized the study, prepared the dataset, performed the analysis, and wrote the manuscript. MER performed the analysis, and wrote the manuscript. NBN conceptualized the study and wrote the manuscript. PDLR conceptualized the study, performed the analysis, wrote the manuscript, and supervised the project. AB conceptualized the study. All authors discussed the results and revised the manuscript.

## Acknowledgments

DM thanks the Swiss National Science Foundation (SNFS) for financial support under grant number P2ELP3_181910. NBN thanks National Health and Medical Research Council of Australia Investigator Grant APP1197021 and Recruitment Grant from Monash University Faculty of Medicine Nursing and Health Sciences with funding from the State Government of Victoria and the Australian Government.

