## Supplementary Material for "Data-driven large-scale genomic analysis reveals an intricate phylogenetic and functional landscape in J-domain proteins"

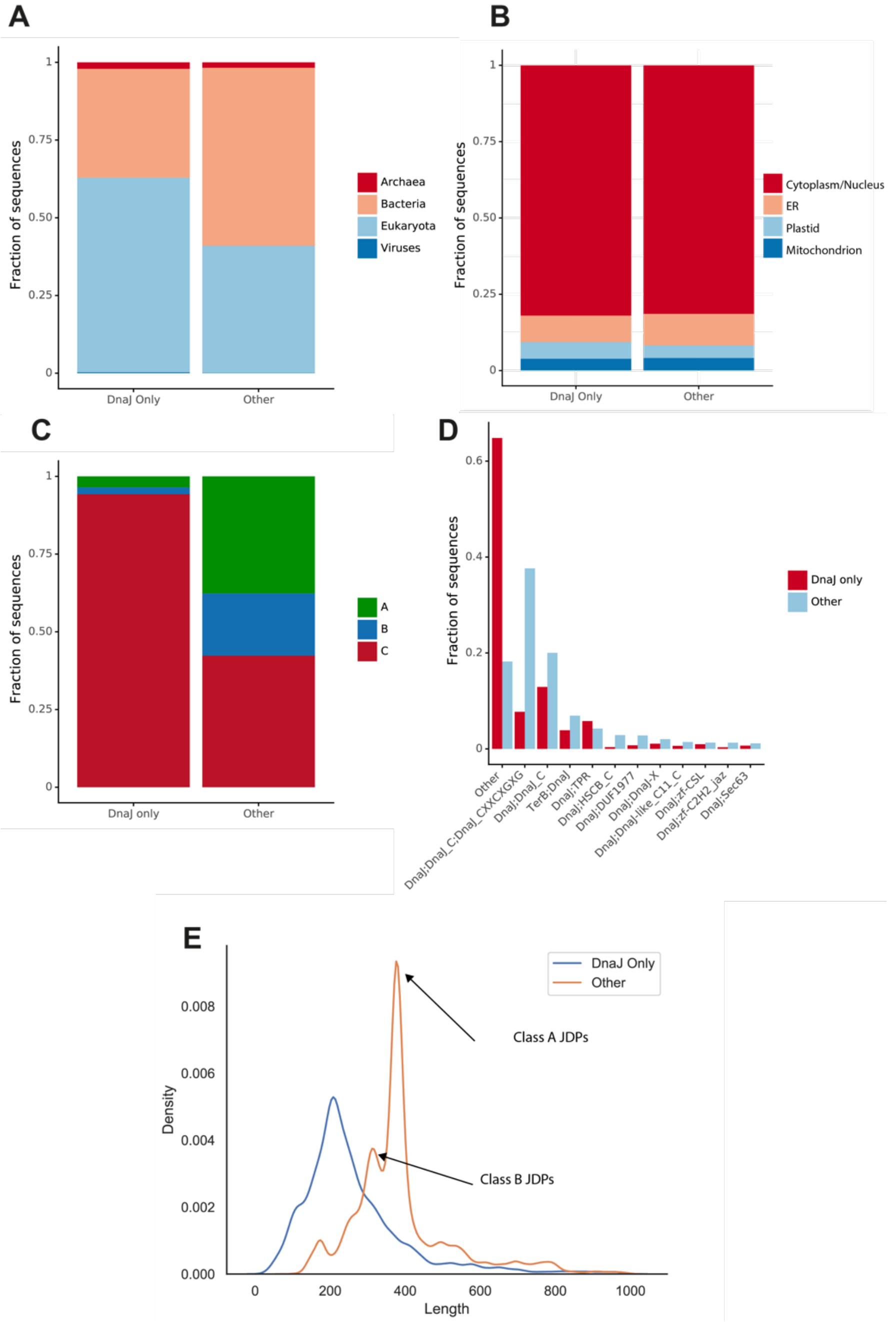


**Supplementary Figure 1. Compositional analysis of J-protein containing only a J-domain. A)** Comparison of the fraction of J-proteins in our dataset belonging to various phylogenetic kingdoms, between JDPs containing only the J-domain ("DnaJ only") and JDPs containing multiple domains ("Other"). **B)** Comparison of the fraction of Eukaryotic J-proteins belonging to subcellular localization (ground truth predicted by TargetP), between JDPs containing only the J-domain ("DnaJ only") and JDPs containing multiple domains ("Other"). **C)** Left: Fraction of predicted ABC J-protein classes, predicted on DnaJ only containing J-proteins ("DnaJ only"). Right: Fraction of ABC J-protein classes, as measured in our dataset on J-proteins containing more than one domain ("Other") **D** Red: Fraction of predicted Archs12 domain architectures, predicted on DnaJ only containing J-proteins ("DnaJ only"). Blue: Fraction of Archs12 domain architectures, as measured in our dataset on J-proteins containing more than one domain ("Other")**). E** Length distributions of J-proteins containing only J-domain (blue) and all other multi-domain J-proteins (orange). The length peaks corresponding to class A and B JDPs are highlighted. For visual clarity, the distributions are only shown for JDPs shorter tha 1000 amino-acids (~98% of the dataset).

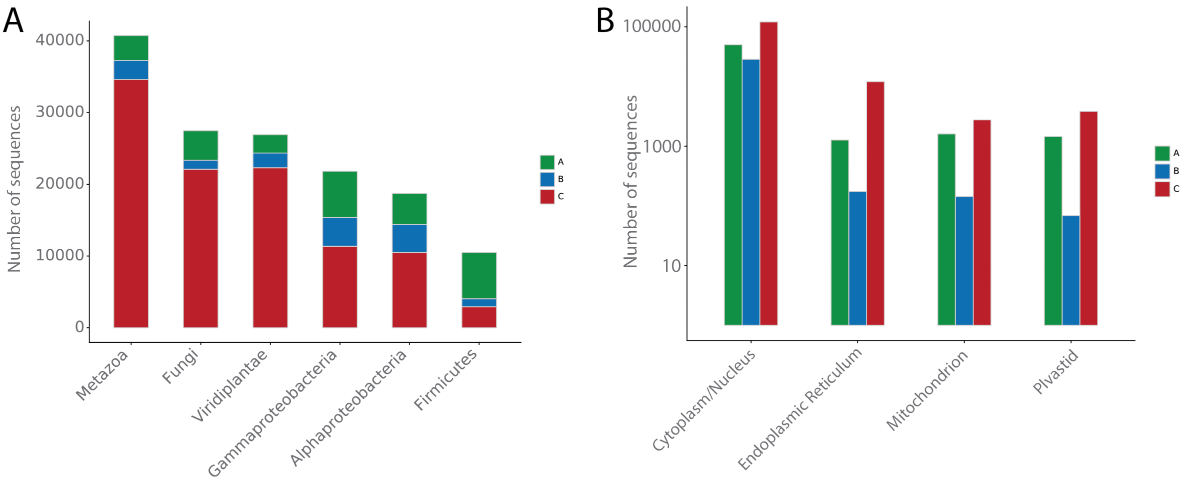


**Supplementary Figure 2. Number of JDP per organism and sub-cellular distribution of JDPs in eukaryotes. A**) Class A, B and C distributions stratified by phylogenetic sub-groups. **B)** Predicted subcellular localization distribution of JDPs in classes A, B and C.

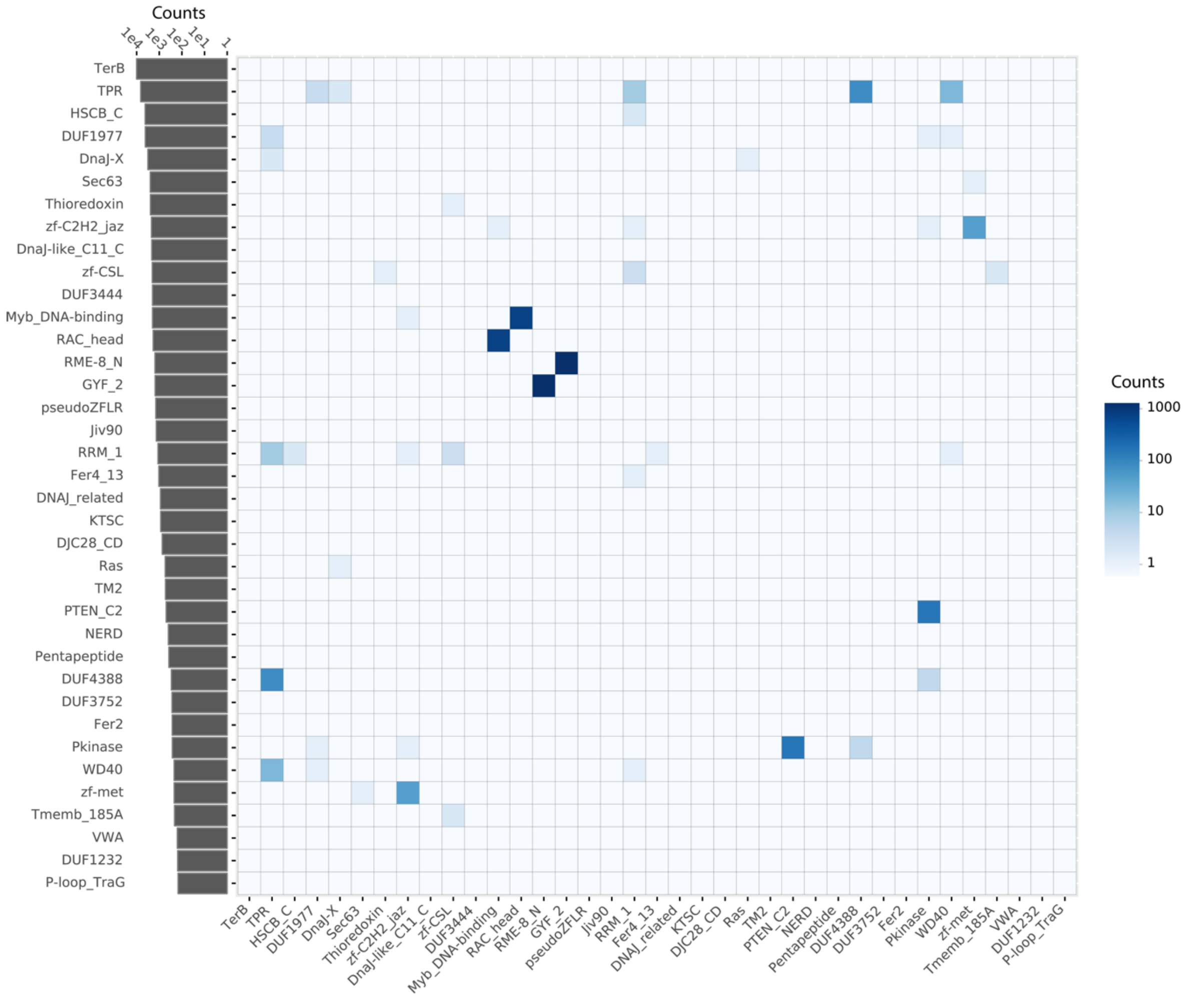


**Supplementary Figure 3. Co-occurrence matrix of the most common domains found in class C J-domain proteins.** The color-scale indicates the number of times a pair-of domain co-occurs in JDPs in our full dataset (ignoring the order of the domain-architectures and domain multiplicity).

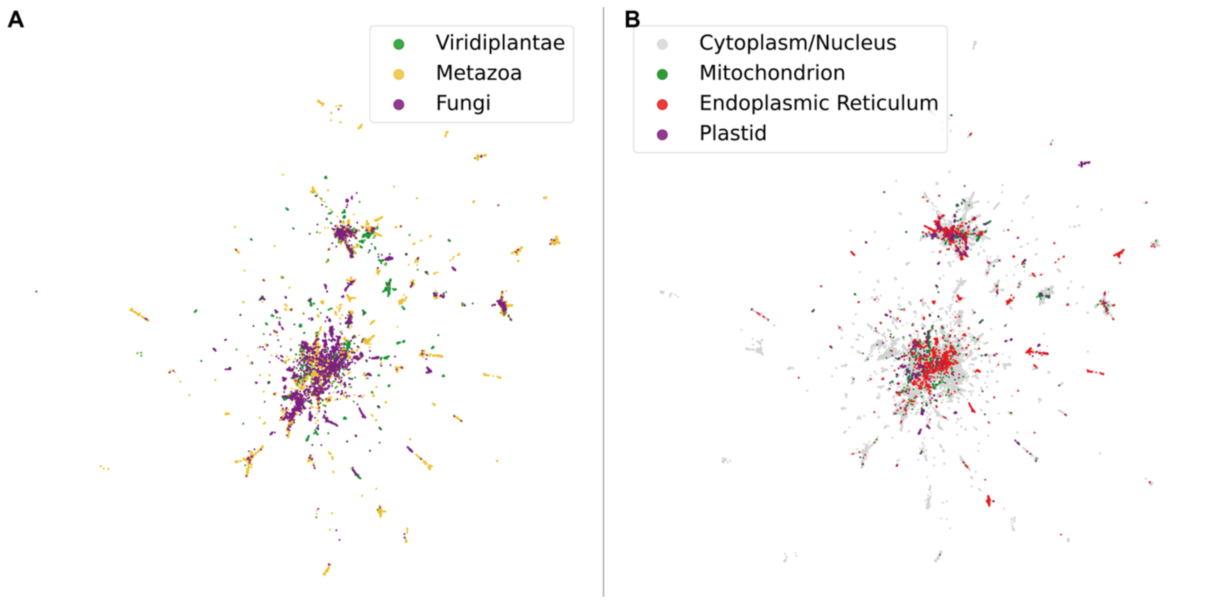


**Supplementary Figure 4. Fine-grained phylogenetic classification and subcellular localization of J-domains.** Shown are Two-dimensional UMAP representations of J-domains as in Fig.3. JDP sequences are projected in two dimensions by a non-linear transformation (UMAP) and colored according to different schemes. **A)** J-domains are identified as belonging to Viridiplantae (green), Metazoa (orange) or Fungi (Purple). **B)** J-domains are identified as being in the Nucleus/Cytoplasm (grey), Mitochondrion (Green), ER (Red) and in plastids (Purple).

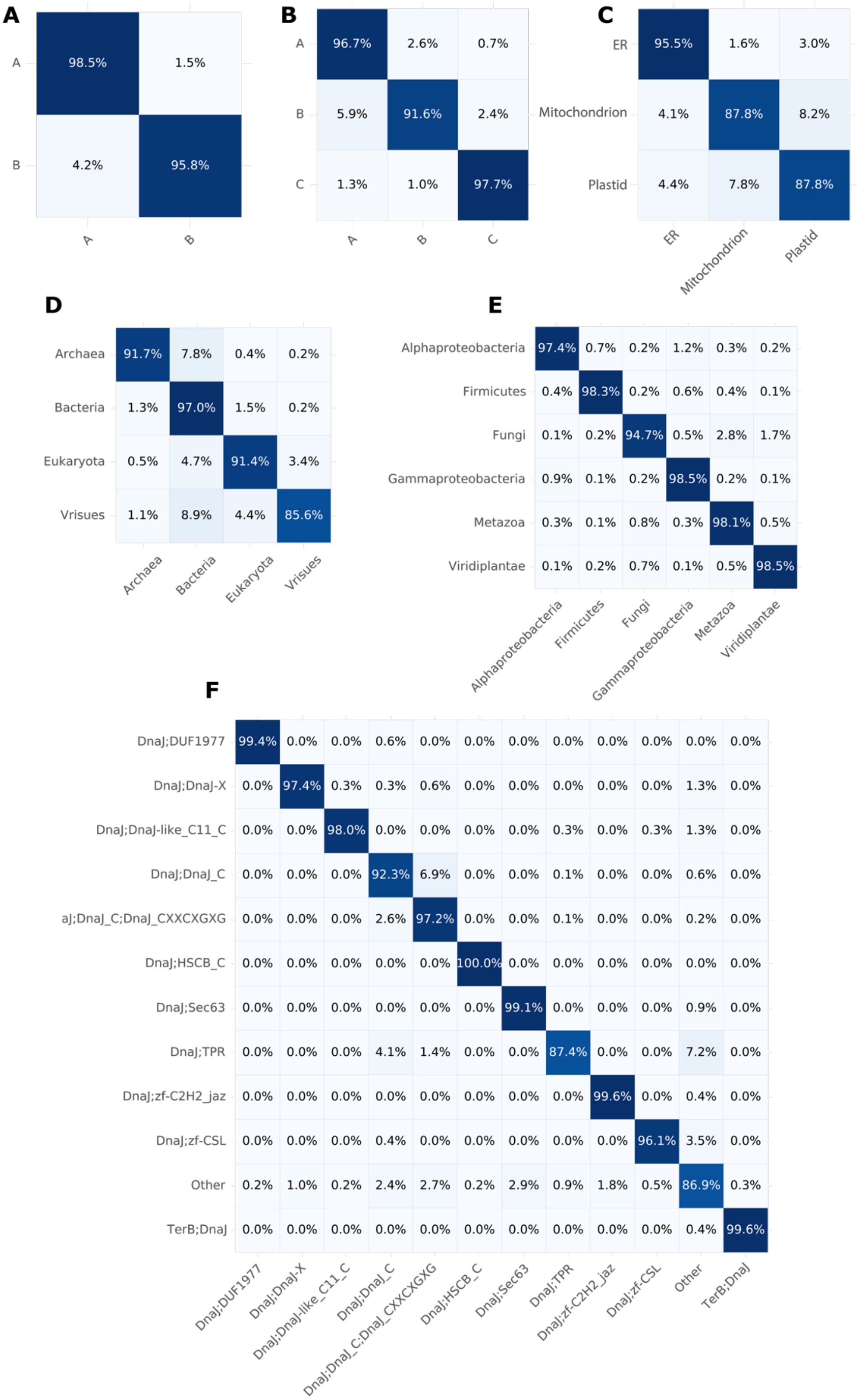


**Supplementary Figure 5. Confusion matrices computed over the test sets for the six multi-label classification tasks discussed in the text.** The matrix row are true label counts vs matrix columns the predicted label counts. **A)** AB task **B)** ABC task **C)** Localization task **D)** Bac/Euk/Arch/Vir task **E)** Phylo task **F)** Top12Archs task.

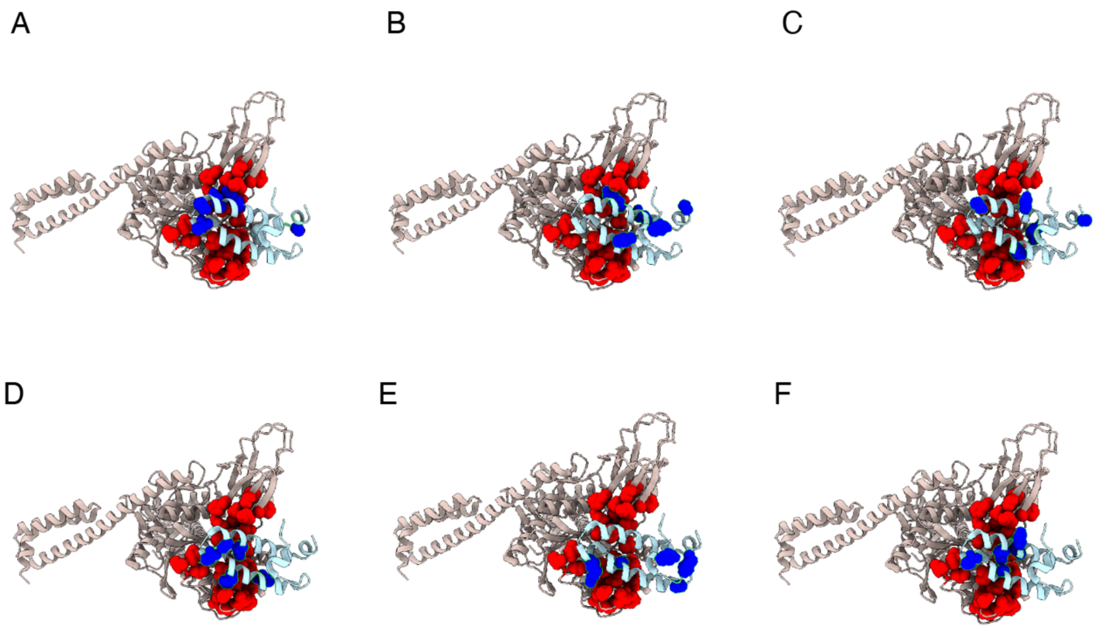


**Supplementary Figure 6. Five most relevant positions for different classification tasks in main Fig.5, mapped on the structure of the complex between DnaK and the J-domain of DnaJ (PDB: 5NRO). A)** A/B. **B)** A/B/C. **C)** Localization **D)** Phylo (as in Supplementary Fig.5B) **E)** Bac/Euk/Arch/Vir **F)** Top12Archs

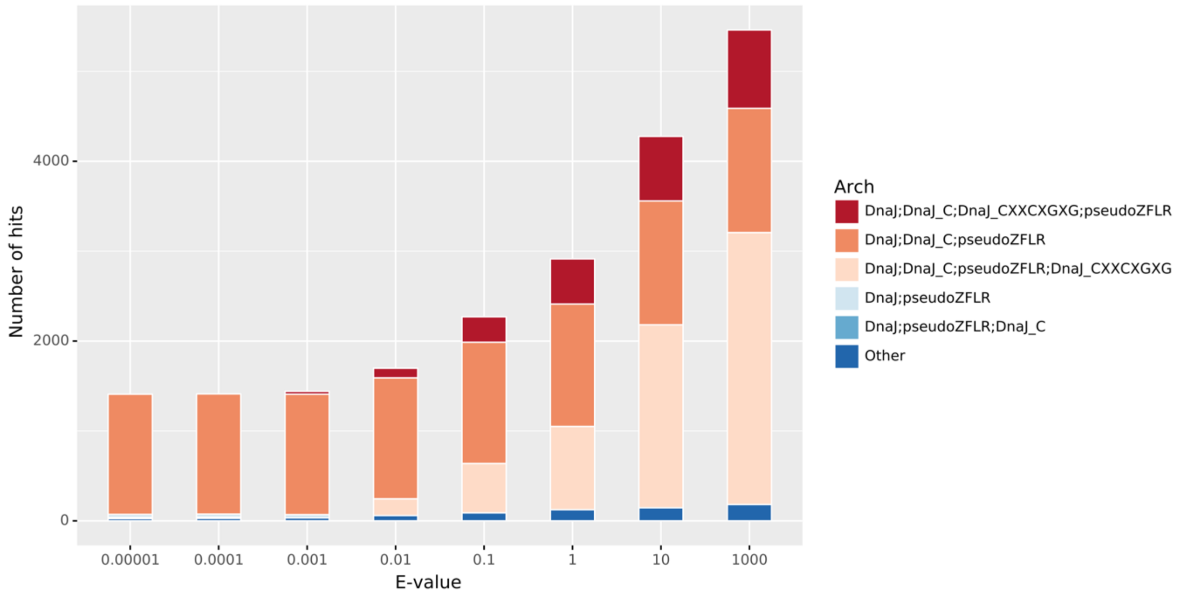


**Supplementary Figure 7. Sensitivity of the identification of pseudoZFLR domains in our JDP dataset.** The x-axis shows the E-value cutoff used with the *hmmsearch* utility. The colors of the stacked bars indicate the various domain architectures in which the pseudo ZF domain is identified with varying selection thresholds. As seen, using more stringent (lower) E-values removes domains containing both the canonical DnaJ_CXXCXGXG and the novel proto-ZF domain (probably corresponding to confused hits, as indeed controlled manually in a small subset of cases), while maintaining a constant number of bona-fide JDPs consisting of DnaJ;DnaJ_C and pseudoZFLR domains.

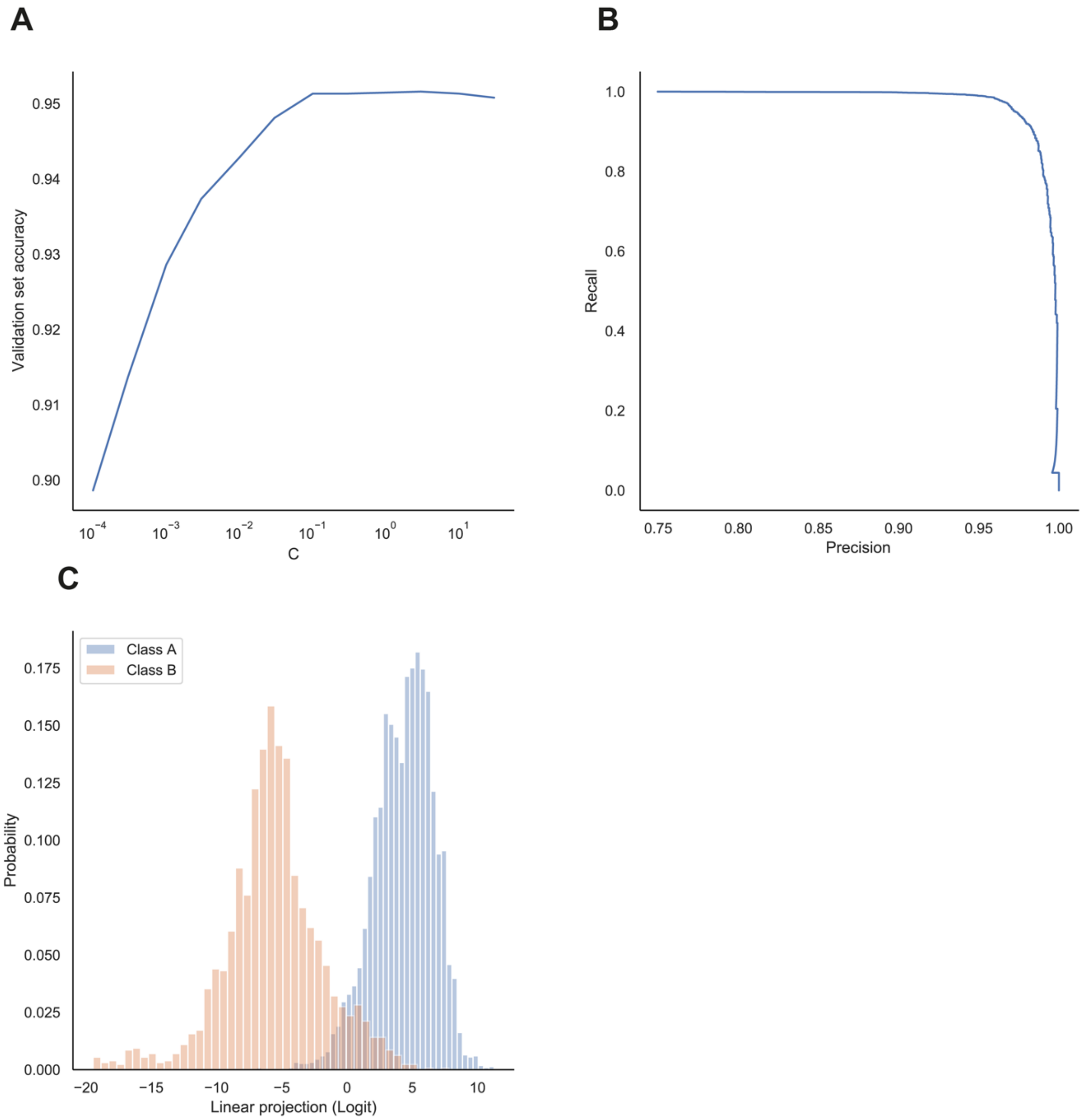


**Supplementary Figure 8. Linear classification of class A vs B JDPs, based on the unaligned G/F regions. A)** Validation set accuracies measured for various regularization strengths C. **B)** Precision-Recall curve computed on the test set for the optimal regularized (C=3.0) logistic regression model. **C)** Distributions of the linear projections (Logits) of class A (blue) and class B (orange) G/F region sequences for the optimal model (C=3.0). The corresponding classification accuracy is ~95%.

**Supplementary Table 1. Classification of the 22 *S. Cerevisiae* (UP000002311 Saccharomyces cerevisiae (strain ATCC 204508 / S288c) (Baker's yeast) (ATCC 204508 / S288c)) JDPs.** Domain architectures grouped into the "Other" class are indicated by (Other) in the third column.

| | *S. Cerevisiae* name | Uniprot | Domain Architecture | A/B/C Probabilities | Arch1-11 Probabilities |
|---|---|---|---|---|---|
| | **Class A** | | | | |
| 1 | MDJ1 | P35191 | DnaJ – DnaJ_C – DnaJ_CXXCXGXG | **1.0** / 0.0 / 0.0 | DnaJ;DnaJ_C;DnaJ_CXXCXGXG **(1.0)** |
| 2 | SCJ1 | P25303 | DnaJ – DnaJ_C – DnaJ_CXXCXGXG | **1.0** / 0.0 / 0.0 | DnaJ;DnaJ_C;DnaJ_CXXCXGXG **(1.0)** |
| 3 | XDJ1 | P39102 | DnaJ – DnaJ_C – DnaJ_CXXCXGXG | 0.13 / 0.0 / **0.87** | DnaJ;DnaJ_C;DnaJ_CXXCXGXG **(0.99)** |
| 4 | YDJ1 | P25491 | DnaJ – DnaJ_C – DnaJ_CXXCXGXG | **0.87** / 0.13 / 0.0 | DnaJ;DnaJ_C;DnaJ_CXXCXGXG **(1.0)** |
| 5 | APJ1 | P53940 | DnaJ – DnaJ_C – DnaJ_CXXCXGXG | **1.0** / 0.0 / 0.0 | DnaJ;DnaJ_C;DnaJ_CXXCXGXG **(1.0)** |
| | **Canonical Class B** | | | | |
| 6 | SIS1 | P25294 | DnaJ – DnaJ_C | 0.0 / **1.0** / 0.0 | DnaJ;DnaJ_C **(0.93)** |
| | **Class C** | | | | |
| 7 | CAJ1 | P39101 | DnaJ – DnaJ-X | 0.0 / 0.0 / **1.0** | DnaJ;DnaJ-X **(1.0)** |
| 8 | JJJ1 | P53863 | DnaJ – zf-C2H2_jaz | 0.0 / 0.0 / **1.0** | DnaJ;zf-C2H2_jaz **(0.99)** |
| **9** | JJJ3 | P47138 | DnaJ – zf-CSL | 0.0 / 0.0 / **1.0** | DnaJ;zf-CSL **(1.0)** |
| 10 | DJP1 | P40564 | DnaJ – DnaJ-X | 0.0 / 0.0 / **1.0** | DnaJ;DnaJ-X **(1.0)** |
| 11 | ZUO1 | P32527 | DnaJ – RAC_head *(Other)* | 0.0 / 0.0 / **1.0** | Other **(0.99)** |
| 12 | JAC1 | P53193 | DnaJ – HSCB_C | 0.0 / 0.0 / **1.0** | DnaJ;HscB_C **(1.0)** |
| 13 | SWA2 | Q06677 | Ubiq-assoc – DnaJ *(Other)* | 0.0 / 0.0 / **1.0** | Other **(1.0)** |
| 14 | HLJ1 | P48353 | DnaJ | 0.0 / 0.0 / **1.0** | DnaJ;DUF1977 **(1.0)** |
| 15 | JEM1 | P40358 | DnaJ | 0.03 / 0.0 / **0.97** | DnaJ;TPR **(0.64)** |
| 16 | JID1 | Q12350 | DnaJ | 0.0 / 0.0 / **1.0** | Other **(0.58)** |
| 17 | ERJ5 | P43613 | DnaJ | 0.05 / 0.0 / **0.95** | DnaJ;DnaJ_C;DnaJ_CXXCXGXG **(0.79)** |
| 18 | JJJ2 | P46997 | DnaJ | 0.0 / 0.0 / **1.0** | Other **(1.0)** |
| 19 | SEC63 | P14906 | DnaJ | 0.0 / 0.0 / **1.0** | DnaJ;Sec63 **(1.0)** |
| 20 | CWC23 | P52868 | DnaJ | 0.0 / 0.0 / **1.0** | Other **(0.61)** |
| 21 | PAM18 | Q07914 | DnaJ | 0.0 / 0.0 / **1.0** | Other **(1.0)** |
| 22 | MDJ2 | P42834 | DnaJ | 0.0 / 0.0 / **1.0** | Other **(1.0)** |

**Supplementary Table 2. Classification of the 9 *H. sapiens* (UP000005640 (Homo sapiens)) class A and canonical class B JDPs.**

| | ***H. Sapiens* name** | **Uniprot** | **Domain Architecture** | **A/B Probabilities** |
|---|---|---|---|---|
| | **Class A** | | | |
| 1 | DnaJA1 | P31689 | DnaJ - DnaJ_C - DnaJ_CXXCXGXG | **1.0** / 0.0 |
| 2 | DnaJA2 | O60884 | DnaJ - DnaJ_C - DnaJ_CXXCXGXG | **0.99** / 0.01 |
| 3 | DnaJA3 | Q96EY1 | DnaJ - DnaJ_C - DnaJ_CXXCXGXG | **1.0** / 0.0 |
| 4 | DnaJA4 | Q8WW22 | DnaJ - DnaJ_C - DnaJ_CXXCXGXG | **1.0** / 0.0 |
| | **Canonical Class B** | | | |
| 5 | DnaJB1 | P25685 | DnaJ - DnaJ_C | 0.0 / **1.0** |
| 6 | DnaJB4 | Q9UDY4 | DnaJ - DnaJ_C | 0.0 / **1.0** |
| 7 | DnaJB5 | O75953 | DnaJ - DnaJ_C | 0.0 / **1.0** |
| 8 | DnaJB11 | Q9UBS4 | DnaJ - DnaJ_C | 0.01 / **0.99** |
| 9 | DnaJB13 | P59910 | DnaJ - DnaJ_C | 0.01 / **0.99** |

**Supplementary Table 3. Classification of the 99 *A. thaliana* (UP000006548 Arabidopsis thaliana (Mouse-ear cress) (cv. Columbia)) JDPs.**

| Gene name | Uniprot | Domain Architecture | A/B/C Probabilities | Arch12 Probabilities |
|---|---|---|---|---|
| **Class A** | | | | |
| ATJ1 | Q38813 | DnaJ - DnaJ_C - DnaJ_CXXCXGXG | **0.8**/0.14/0.06 | DnaJ;DnaJ_C;DnaJ_CXXCXGXG (**0.94**) |
| ATJ2 | P42825 | DnaJ - DnaJ_C - DnaJ_CXXCXGXG | **0.9**/0.1/0.01 | DnaJ;DnaJ_C;DnaJ_CXXCXGXG (**0.98**) |
| ATJ3 | Q94AW8 | DnaJ - DnaJ_C - DnaJ_CXXCXGXG | **0.86**/0.13/0.01 | DnaJ;DnaJ_C;DnaJ_CXXCXGXG (**1.0**) |
| DJA5 | Q940V1 | DnaJ - DnaJ_C - DnaJ_CXXCXGXG | **1.0**/0.0/0.0 | DnaJ;DnaJ_C;DnaJ_CXXCXGXG (**0.99**) |
| DJA6 | Q9SJZ7 | DnaJ - DnaJ_C - DnaJ_CXXCXGXG | **1.0**/0.0/0.0 | DnaJ;DnaJ_C;DnaJ_CXXCXGXG (**1.0**) |
| GFA2 | Q8GWW8 | DnaJ - DnaJ_C - DnaJ_CXXCXGXG | **0.73**/0.22/0.05 | DnaJ;DnaJ_C;DnaJ_CXXCXGXG (**0.93**) |
| DJA7 | Q0WN54 | DnaJ - DnaJ_C - DnaJ_CXXCXGXG | **1.0**/0.0/0.0 | DnaJ;DnaJ_C;DnaJ_CXXCXGXG (**0.94**) |
| DJA4 | Q058J9 | DnaJ - DnaJ_C - DnaJ_CXXCXGXG | **0.92**/0.0/0.8 | DnaJ;DnaJ_C;DnaJ_CXXCXGXG (**0.73**) |
| **Canonical Class B** | | | | |
| ERDJ3B | Q9LZK5 | DnaJ - DnaJ_C - pseudoZFLR | 0.0/0.0/**1.0** | Other (**0.96**) |
| F20O9.160 | O49457 | DnaJ - DnaJ_C | 0.0/**1.0**/0.0 | DnaJ;DnaJ_C (**1.0**) |
| F23H11.4 | Q9XIF5 | DnaJ - DnaJ_C | 0.0/**1.0**/0.0 | DnaJ;DnaJ_C (**0.99**) |
| T13C7.15 | Q9SIL3 | DnaJ - DnaJ_C | 0.0/**1.0**/0.0 | DnaJ;DnaJ_C (**1.0**) |
| T14C9.70 | F4JY55 | DnaJ - DnaJ_C | 0.0/**1.0**/0.0 | DnaJ;DnaJ_C (**1.0**) |
| T16O11.15 | Q9SR91 | DnaJ - DnaJ_C | 0.0/**1.0**/0.0 | DnaJ;DnaJ_C (**1.0**) |
| AT1G10350 | Q9SY77 | DnaJ;DnaJ_C | 0.0/**1.0**/0.0 | DnaJ;DnaJ_C (**1.0**) |
| T17F15.190 | Q9SU57 | DnaJ;DnaJ_C | 0.0/**1.0**/0.0 | DnaJ;DnaJ_C (**1.0**) |
| T10O8.100 | F4K9C8 | DnaJ;DnaJ_C | 0.0/**1.0**/0.0 | DnaJ;DnaJ_C (**0.99**) |
| **Class C** | | | | |
| ATJ10 | Q8GYX8 | DnaJ - DnaJ-X | 0.0/0.0/**1.0** | DnaJ;DnaJ-X (**1.0**) |
| ATJ11 | Q9FYB5 | DnaJ | 0.0/0.0/**1.0** | Other (**0.97**) |
| ATJ13 | Q39079 | DnaJ - DnaJ-like_C11_C | 0.0/0.0/**1.0** | DnaJ;DnaJ-like_C11_C (**1.0**) |
| ATJ15 | Q9ZSY2 | DnaJ | 0.2/0.04/**0.76** | Other (**0.92**) |
| ATJ16 | Q8VXV4 | DnaJ | 0.0/**0.66**/0.34 | DnaJ;DnaJ_C (**0.5**) |
| ATJ20 | Q9SDN0 | DnaJ | 0.0/0.0/**1.0** | Other (**0.73**) |
| ATJ39 | Q6XL73 | DnaJ | 0.0/0.07/**0.93** | DnaJ;DnaJ_C (**0.58**) |
| ATJ49 | Q9FH28 | DnaJ - DUF1977 | 0.0/0.0/**1.0** | DnaJ;DUF1977 (**1.0**) |
| ATJ6 | Q9FL54 | DnaJ | 0.0/0.01/**0.99** | Other (**0.57**) |
| ATJ72 | Q0WTI8 | DnaJ | 0.0/0.0/**1.0** | DnaJ;DnaJ_C;DnaJ_CXXCXGXG (**0.77**) |
| ATJ8 | Q9SAG8 | DnaJ | 0.0/0.0/**1.0** | DnaJ;DnaJ_C (**0.61**) |
| A_TM021B04.9 | O04648 | DnaJ - DUF3444 - DUF3444 - DDE_Tnp_4 - Myb_DNA-bind_3 | 0.0/0.0/**1.0** | Other (**1.0**) |
| C50 | Q8GUN6 | DnaJ | 0.0/0.0/**1.0** | Other (**0.98**) |
| DJC66 | Q9LTT7 | DnaJ | 0.0/0.0/**1.0** | DnaJ;DnaJ_C (**0.39**) |
| DJC72 | Q8L763 | DnaJ | 0.0/0.0/**1.0** | Other (**1.0**) |
| DJC76 | Q9FMX6 | DnaJ - Fer4_13 | 0.0/0.0/**1.0** | Other (**1.0**) |
| DJC77 | Q9SJI1 | DnaJ - Fer4_13 | 0.0/0.0/**1.0** | Other (**1.0**) |
| ERDJ2A | Q0WT48 | DnaJ - Sec63 | 0.0/0.0/**1.0** | DnaJ;Sec63 (**1.0**) |
| ERDJ2B | F4JIN3 | DnaJ - Sec63 | 0.0/0.0/**1.0** | DnaJ;Sec63 (**1.0**) |
| ERDJ3A | Q9SR96 | DnaJ | 0.19/0.0/**0.81** | DnaJ;DnaJ_C (**0.62**) |
| F13B15.22 | Q9SLA7 | DnaJ - DUF3444 | 0.0/0.0/**1.0** | Other (**1.0**) |
| F14G9.9 | Q8L7R1 | DnaJ | 0.0/0.0/**1.0** | DnaJ;DnaJ_C (**0.89**) |
| F19K19.3 | Q9FX81 | DnaJ - Jiv90 | 0.0/0.0/**1.0** | Other (**0.99**) |
| F24J7.150 | A0A1P8B3S0 | DnaJ | 0.0/0.0/**1.0** | Other (**1.0**) |
| F24P17.19 | Q9SQT7 | DnaJ - DUF3444 | 0.0/0.0/**1.0** | Other (**1.0**) |
| F28J15.2 | Q9LH49 | DnaJ | 0.0/0.0/**1.0** | Other (**0.68**) |
| F28P5.5 | F4IBN6 | DnaJ | 0.0/0.0/**1.0** | Other (**0.9**) |
| F3K23.27 | A0A1P8B0U4 | DnaJ - DnaJ-X | 0.0/0.0/**1.0** | DnaJ;DnaJ-X (**1.0**) |
| F5G3.13 | Q9S7L6 | DnaJ - DUF3444 | 0.0/0.0/**1.0** | Other (**1.0**) |
| GRV2 | F4IVL6 | RME-8_N - RME-8_N - RME-8_N - GYF_2 - DnaJ | 0.0/0.0/**1.0** | Other (**1.0**) |
| JJJ1 | Q9C911 | DnaJ - zf-C2H2_jaz | 0.0/0.0/**1.0** | DnaJ;zf-C2H2_jaz (**1.0**) |
| K12B20.210 | F4K8L9 | DnaJ | 0.0/0.0/**1.0** | Other (**1.0**) |
| K12B20.23 | F4K8L8 | DnaJ | 0.0/0.0/**1.0** | Other (**1.0**) |
| MFH8.9 | Q6NMG4 | DnaJ - DUF3444 | 0.0/0.0/**1.0** | Other (**1.0**) |
| MNJ8.20 | Q9FHS7 | DnaJ | 0.0/0.0/**1.0** | Other (**1.0**) |
| MQB2.10 | F4K7T4 | DnaJ | 0.0/0.0/**1.0** | Other (**1.0**) |
| MRG7.10 | Q9FK56 | DnaJ | 0.0/0.0/**1.0** | Other (**0.91**) |
| MTG13.10 | A0A1P8BDW0 | DnaJ | 0.0/0.0/**1.0** | Other (**0.93**) |
| MTH12.18 | Q9FNA0 | DnaJ | 0.0/0.0/**1.0** | Other (**0.77**) |
| At3g11450 | F4J6A8 | DnaJ - Myb_DNA-binding | 0.0/0.0/**1.0** | Other (**1.0**) |
| P58IPK | Q9LYW9 | DnaJ - TPR | 0.0/0.0/**1.0** | DnaJ;TPR (**1.0**) |
| T22I11.9 | F4HWD5 | DnaJ - DnaJ-X | 0.0/0.0/**1.0** | DnaJ;DnaJ-X (**1.0**) |
| T25O11.9 | Q9FG44 | DnaJ | 0.0/0.0/**1.0** | Other (**1.0**) |
| T6D20.3 | P93739 | DnaJ | 0.0/0.0/**1.0** | Other (**0.77**) |
| T8F5.5 | Q0WVU7 | DnaJ - DUF3752 | 0.0/0.0/**1.0** | Other (**1.0**) |
| T8O11.12 | Q9ZU99 | DnaJ | 0.0/0.0/**1.0** | Other (**1.0**) |
| T9J14.7 | A0A1I9LPC5 | DnaJ - DUF3444 - DUF3444 | 0.0/0.0/**1.0** | Other (**1.0**) |
| TPR15 | A0A1P8B1G1 | DnaJ | 0.0/0.0/**1.0** | DnaJ;TPR (**1.0**) |
| TPR16 | F4K0Y5 | DnaJ - TPR | 0.0/0.0/**1.0** | DnaJ;TPR (**1.0**) |
| ndhT | Q9SMS0 | DnaJ | 0.0/0.0/**1.0** | Other (**0.99**) |
| At1g09260 | O80481 | DnaJ | 0.0/0.0/**1.0** | Other (**0.98**) |
| AT1G18700 | Q8VYU5 | DnaJ | 0.0/0.0/**1.0** | Other (**1.0**) |
| At1g62970 | Q0WQM3 | DnaJ | 0.0/0.0/**1.0** | Other (**1.0**) |
| At1g71000 | F4I8D6 | DnaJ | 0.0/0.0/**1.0** | DnaJ;DnaJ_C;DnaJ_CXXCXGXG (**0.6**) |
| AT1G72416 | Q1G3W9 | DnaJ | 0.0/0.0/**1.0** | Other (**0.98**) |
| At1g77020 | F4I5J7 | DnaJ;DnaJ-X | 0.0/0.0/**1.0** | DnaJ;DnaJ-X (**1.0**) |

| | | | | |
|---|---|---|---|---|
| DJC65 | Q9SH08 | DnaJ | 0.0/0.0/**1.0** | Other (**1.0**) |
| YUP8H12R.35 | F4IDH6 | DnaJ;Jiv90 | 0.0/0.0/**1.0** | Other (**1.0**) |
| DJC24 | O48828 | DnaJ | 0.0/0.0/**1.0** | DnaJ;DnaJ_C;DnaJ_CXXCXGXG (**0.74**) |
| AT2G33735 | Q8RYC5 | DnaJ | 0.0/0.0/**1.0** | Other (**0.54**) |
| At2g35540 | F4IKR5 | DnaJ;DUF3444 | 0.0/0.0/**1.0** | Other (**1.0**) |
| At2g42065 | A0A1P8AYV4 | DnaJ | 0.0/0.0/**1.0** | Other (**0.78**) |
| DJC82 | F4J794 | DnaJ | 0.0/0.0/**1.0** | Other (**1.0**) |
| At3g06778 | F4JC50 | DnaJ | 0.0/0.0/**1.0** | Other (**1.0**) |
| At3g14200 | Q9LJG5 | DnaJ | 0.0/0.0/**1.0** | Other (**0.75**) |
| F28O9.190 | Q9M2L3 | DnaJ;DUF1977 | 0.0/0.0/**1.0** | DnaJ;DUF1977 (**1.0**) |
| At3g58020 | Q8RXJ5 | DnaJ | 0.0/0.0/**1.0** | Other (**1.0**) |
| At3g62190 | C0Z334 | DnaJ | 0.0/0.0/**1.0** | Other (**0.99**) |
| At4g07990 | Q8L814 | DnaJ | 0.0/0.0/**1.0** | DnaJ;TPR (**0.78**) |
| T9A4.1 | O82623 | DnaJ;zf-CSL | 0.0/0.0/**1.0** | DnaJ;zf-CSL (**1.0**) |
| AT4G19570 | Q84TH2 | DnaJ | 0.0/0.0/**1.0** | Other (**1.0**) |
| At4g19580 | F4JT89 | DnaJ;WCCH | 0.0/0.0/**1.0** | Other (**1.0**) |
| At4g37480 | F4JS27 | DnaJ | 0.0/0.0/**1.0** | Other (**0.79**) |
| At4g39150 | Q9T024 | DnaJ;DnaJ-X | 0.0/0.0/**1.0** | DnaJ;DnaJ-X (**1.0**) |
| MJJ3.16 | Q9FFK2 | DnaJ | 0.0/0.0/**1.0** | DnaJ;DUF1977 (**1.0**) |
| AT5G06110 | Q9LHS5 | DnaJ;Myb_DNA-binding | 0.0/0.0/**1.0** | Other (**1.0**) |
| At5g09540 | Q9LXB9 | DnaJ | 0.0/0.0/**1.0** | Other (**1.0**) |
| AT5G18750 | Q3E9D9 | DnaJ;DUF3444;DUF3444 | 0.0/0.0/**1.0** | Other (**0.65**) |
| At5g22080 | Q9C580 | DnaJ | 0.0/0.0/**1.0** | Other (**0.97**) |
| MQM1.14 | Q8L7M3 | DnaJ;RRM_1 | 0.0/0.0/**1.0** | Other (**1.0**) |
| AT5G49580 | Q9FGY8 | DnaJ;Jiv90 | 0.0/0.0/**1.0** | Other (**1.0**) |
| EIP9 | Q9FMF3 | DnaJ | 0.0/0.0/**1.0** | Other (**1.0**) |

**Supplementary Table 4. Classification of the 31 *C. elegans* (UP000001940 Caenorhabditis elegans (Bristol N2)) JDPs.**

| Gene name | Uniprot | Domains | A/B/C Probabilities | Arch12 Probabilities |
|---|---|---|---|---|
| **Class A** | | | | |
| dnj-10 | Q8TA83 | DnaJ - DnaJ_C - DnaJ_CXXCXGXG | **1.0**/0.0/0.0 | DnaJ;DnaJ_C;DnaJ_CXXCXGXG (**0.98**) |
| dnj-12 | O45502 | DnaJ - DnaJ_C - DnaJ_CXXCXGXG | **0.7**/0.29/0.0 | DnaJ;DnaJ_C (**0.61**) |
| dnj-19 | O16303 | DnaJ - DnaJ_C - DnaJ_CXXCXGXG | **0.84**/0.06/0.1 | DnaJ;DnaJ_C;DnaJ_CXXCXGXG (**0.99**) |
| **Canonical Class B** | | | | |
| dnj-13 | Q20774 | DnaJ - DnaJ_C | 0.0/**1.0**/0.0 | DnaJ;DnaJ_C (**1.0**) |
| dnj-20 | Q8MPX3 | DnaJ - DnaJ_C - pseudoZFLR | 0.02/0.08/**0.9** | Other (**0.98**) |
| **Class C** | | | | |
| CELE_K07F5.16 | Q7YWX3 | DnaJ | 0.0/0.0/**1.0** | Other (**1.0**) |
| F54F2.9 | P34454 | DnaJ - Myb_DNA-binding | 0.0/0.0/**1.0** | Other (**1.0**) |
| dnj-1 | Q17438 | DnaJ - DUF1977 | 0.0/0.0/**1.0** | DnaJ;DUF1977 (**1.0**) |
| dnj-11 | Q94216 | DnaJ - RAC_head - Myb_DNA-binding | 0.0/0.0/**1.0** | Other (**1.0**) |
| dnj-14 | Q21162 | DnaJ | 0.0/0.0/**1.0** | Other (**1.0**) |
| dnj-15 | Q21324 | DnaJ - HSCB_C | 0.0/0.0/**1.0** | DnaJ;HSCB_C (**1.0**) |
| dnj-16 | Q22028 | DnaJ | 0.01/0.28/**0.7** | DnaJ;DnaJ_C;DnaJ_CXXCXGXG (**0.93**) |
| dnj-17 | O62360 | DnaJ - zf-C2H2_jaz | 0.0/0.0/**1.0** | DnaJ;zf-C2H2_jaz (**1.0**) |
| dnj-18 | Q22138 | DnaJ | 0.0/0.0/**1.0** | Other (**0.99**) |
| dnj-2 | Q17433 | DnaJ | 0.0/0.0/**1.0** | Other (**0.99**) |
| dnj-22 | O17002 | DnaJ | 0.0/0.0/**1.0** | DnaJ;DnaJ_C (**0.87**) |
| dnj-23 | Q22751 | DnaJ | 0.0/0.0/**1.0** | Other (**0.57**) |
| dnj-24 | Q9TY07 | DnaJ | 0.01/0.0/**0.99** | Other (**0.53**) |
| dnj-25 | G5EDJ6 | DnaJ | 0.0/0.0/**1.0** | Other (**1.0**) |
| dnj-26 | Q9U2L4 | DnaJ | 0.0/0.0/**1.0** | Other (**1.0**) |
| dnj-27 | Q9XWE1 | DnaJ - Thioredoxin - Thioredoxin - Thioredoxin - Thioredoxin | 0.0/0.0/**1.0** | Other (**1.0**) |
| dnj-28 | Q9N3E0 | DnaJ - TPR | 0.0/0.0/**1.0** | DnaJ;TPR (**1.0**) |
| dnj-29 | Q9U1W0 | DnaJ - Sec63 | 0.0/0.0/**1.0** | DnaJ;Sec63 (**1.0**) |
| dnj-3 | Q93171 | DnaJ | 0.0/0.0/**1.0** | DnaJ;zf-CSL (**0.67**) |
| dnj-30 | Q95Y44 | DnaJ | 0.0/0.0/**1.0** | Other (**1.0**) |
| dnj-4 | P91014 | DnaJ | 0.0/0.0/**1.0** | Other (**1.0**) |
| dnj-7 | P91189 | DnaJ - TPR | 0.0/0.0/**1.0** | DnaJ;TPR (**1.0**) |
| dnj-8 | Q95QQ1 | DnaJ - Thioredoxin | 0.01/0.0/**0.99** | DnaJ;DnaJ_C (**0.99**) |
| dnj-9 | P91243 | DnaJ - DnaJ-like_C11_C | 0.0/0.0/**1.0** | DnaJ;DnaJ-like_C11_C (**1.0**) |
| rme-8 | G5ED36 | RME-8_N - GYF_2 - DnaJ | 0.0/0.0/**1.0** | Other (**1.0**) |
| Dnj-5 | Q09446 | DnaJ – Jiv90 | 0.0/0.0/**1.0** | DnaJ;TPR (**0.98**) |

**Supplementary Table 5. Classification of the 45 *D. rerio* (UP000814640 Danio rerio (Zebrafish) (Brachydanio rerio) strain Tuebingen) JDPs.**

| Gene name | Uniprot | Domains | A/B/C Probabilities | Arch12 Probabilities |
|---|---|---|---|---|
| **Class A** | | | | |
| dnaja1 | F1QTK7 | DnaJ - DnaJ_C - DnaJ_CXXCXGXG | **1.0**/0.0/0.0 | DnaJ;DnaJ_C;DnaJ_CXXCXGXG (**0.99**) |
| dnaja2b | A5WVI9 | DnaJ - DnaJ_C - DnaJ_CXXCXGXG | **1.0**/0.0/0.0 | DnaJ;DnaJ_C;DnaJ_CXXCXGXG (**1.0**) |
| dnaja3b | B0S5B7 | DnaJ - DnaJ_C - DnaJ_CXXCXGXG | **1.0**/0.0/0.0 | DnaJ;DnaJ_C;DnaJ_CXXCXGXG (**0.99**) |
| **Canonical Class B** | | | | |
| dnajb5 | A5WUU0 | DnaJ - DnaJ_C | 0.0/**0.99**/0.01 | DnaJ;DnaJ_C (**0.99**) |
| dnajb11 | Q6NYZ0 | DnaJ - DnaJ_C - pseudoZFLR | 0.0/0.04/**0.96** | Other (**0.99**) |
| zgc:122979 | F1R1Q6 | DnaJ - DnaJ_C | 0.0/**0.98**/0.02 | DnaJ;DnaJ_C (**0.67**) |
| dnjab1a | A0A0R4IYH3 | DnaJ - DnaJ_C | 0.0/0.0/**1.0** | DnaJ;DnaJ_C (**1.0**) |
| dnjab1b | B8JM56 | DnaJ - DnaJ_C | 0.0/0.0/**1.0** | DnaJ;DnaJ_C (**1.0**) |
| **Class C** | | | | |
| dnajb12a | Q7SXR2 | DnaJ - DUF1977 | 0.02/0.0/**0.98** | DnaJ;DUF1977 (**1.0**) |
| dnajb12b | F1QW75 | DnaJ - DUF1977 | 0.0/0.0/**1.0** | DnaJ;DUF1977 (**1.0**) |
| dnajb14 | A0JMH5 | DnaJ - DUF1977 | 0.0/0.0/**1.0** | DnaJ;DUF1977 (**1.0**) |
| dnajb2 | F1R1B0 | DnaJ | 0.4/0.12/**0.48** | DnaJ;DnaJ_C (**0.75**) |
| dnajb6a | Q6DHR2 | DnaJ | **0.58**/0.15/0.27 | DnaJ;DnaJ_C;DnaJ_CXXCXGXG (**0.73**) |
| dnajb6b | Q1LY18 | DnaJ | 0.0/**0.76**/0.24 | DnaJ;DnaJ_C (**0.93**) |
| dnajb9a | F1QB06 | DnaJ | **0.52**/0.0/0.48 | DnaJ;DnaJ_C;DnaJ_CXXCXGXG (**0.6**) |
| dnajb9b | A5WWB9 | DnaJ | **0.75**/0.0/0.25 | DnaJ;DnaJ_C;DnaJ_CXXCXGXG (**0.92**) |
| dnajc1 | B0UXV7 | DnaJ - Myb_DNA-binding | 0.0/0.0/**1.0** | Other (**1.0**) |
| dnajc10 | A4IG47 | DnaJ - Thioredoxin - Thioredoxin - Thioredoxin - Thioredoxin | 0.0/0.0/**1.0** | Other (**1.0**) |
| dnajc11a | F1QWW6 | DnaJ - DnaJ-like_C11_C | 0.0/0.0/**1.0** | DnaJ;DnaJ-like_C11_C (**1.0**) |
| dnajc11b | H9GYR4 | DnaJ - DnaJ-like_C11_C | 0.0/0.0/**1.0** | DnaJ;DnaJ-like_C11_C (**1.0**) |
| dnajc12 | E7F5S9 | DnaJ | 0.0/0.0/**1.0** | Other (**1.0**) |
| dnajc14 | A0A140LGQ6 | DnaJ - Jiv90 | 0.0/0.0/**1.0** | Other (**0.96**) |
| dnajc16l | F1Q755 | DnaJ - Thioredoxin | 0.0/0.01/**0.99** | DnaJ;DnaJ_C (**0.92**) |
| dnajc17 | A0A0R4ILG8 | DnaJ - RRM_1 | 0.0/0.0/**1.0** | Other (**1.0**) |
| dnajc19 | Q6PBT7 | DnaJ | 0.0/0.0/**1.0** | Other (**1.0**) |
| dnajc2 | Q6NWJ4 | DnaJ - RAC_head - Myb_DNA-binding - Myb_DNA-binding | 0.0/0.0/**1.0** | Other (**1.0**) |
| dnajc21 | Q6PGY5 | DnaJ - zf-C2H2_jaz - zf-C2H2_2 | 0.0/0.0/**1.0** | DnaJ;zf-C2H2_jaz (**1.0**) |
| dnajc22 | E7F2F5 | TM2 - DnaJ | 0.0/0.0/**1.0** | Other (**1.0**) |
| dnajc24 | Q7SY41 | DnaJ - zf-CSL | 0.0/0.0/**1.0** | DnaJ;zf-CSL (**1.0**) |
| dnajc25 | F1RA77 | DnaJ | 0.0/0.0/**1.0** | Other (**1.0**) |
| dnajc27 | Q6IMK3 | Ras - DnaJ | 0.0/0.0/**1.0** | Other (**1.0**) |
| dnajc28 | B0R066 | DnaJ - DJC28_CD | 0.0/0.0/**1.0** | Other (**1.0**) |
| dnajc30b | E7F8P1 | DnaJ | 0.0/0.0/**1.0** | DnaJ;DnaJ-X (**1.0**) |
| dnajc3a | Q6P0U6 | DnaJ - TPR | 0.0/0.0/**1.0** | DnaJ;TPR (**1.0**) |
| dnajc5aa | Q6DGJ0 | DnaJ | 0.01/0.0/**0.99** | Other (**0.97**) |
| dnajc5ab | E7FF34 | DnaJ | 0.01/0.01/**0.99** | Other (**1.0**) |
| dnajc5b | E7F596 | DnaJ | 0.01/0.0/**0.99** | Other (**1.0**) |
| dnajc5ga | Q7ZW85 | DnaJ | 0.0/0.07/**0.93** | Other (**0.96**) |
| dnajc5gb | F1RD67 | DnaJ | 0.02/0.01/**0.96** | Other (**0.61**) |
| dnajc7 | F1QNW8 | DnaJ - TPR | 0.01/0.01/**0.99** | DnaJ;TPR (**1.0**) |
| dnajc8 | Q1ED27 | DnaJ | 0.0/0.0/**1.0** | Other (**0.53**) |
| dnajc9 | Q6DH49 | DnaJ | 0.0/0.0/**1.0** | Other (**1.0**) |
| Hscb | B0V1J6 | DnaJ - HSCB_C | 0.0/0.0/**1.0** | DnaJ;HSCB_C (**1.0**) |
| sec63 | F1QGI2 | DnaJ - Sec63 - Sec63 | 0.0/0.0/**1.0** | DnaJ;Sec63 (**0.85**) |
| zgc:152986 | F1QY22 | DnaJ | 0.01/0.0/**0.99** | Other (**1.0**) |

**Supplementary Table 6. Classification of the 45 *D. melanogaster* (UP000000803 Drosophila melanogaster (Fruit fly) (Berkeley)) JDPs.**

| Gene name | Uniprot | Domains | A/B/C Probabilities | Arch12 Probabilities |
|---|---|---|---|---|
| **Class A** | | | | |
| Droj2 | Q9VFV9 | DnaJ - DnaJ_C - DnaJ_CXXCXGXG | **0.63**/0.36/0.01 | DnaJ;DnaJ_C;DnaJ_CXXCXGXG (**0.9**) |
| DnaJ-H | Q9VK35 | DnaJ - DnaJ_C - DnaJ_CXXCXGXG | **0.99**/0.0/0.0 | DnaJ;DnaJ_C;DnaJ_CXXCXGXG (**0.98**) |
| l(2)tid | Q27237 | DnaJ - DnaJ_C - DnaJ_CXXCXGXG | **0.98**/0.01/0.01 | DnaJ;DnaJ_C;DnaJ_CXXCXGXG (**0.95**) |
| **Canonical Class B** | | | | |
| Dmel\CG2887 | Q9W2U5 | DnaJ - DnaJ_C | 0.0/0.03/**0.97** | DnaJ;DnaJ_C (**0.93**) |
| Dmel\CG5001 | Q9VPY9 | DnaJ - DnaJ_C | 0.0/**1.0**/0.0 | DnaJ;DnaJ_C (**0.99**) |
| Dmel\CG7387 | Q8SZX1 | DnaJ - DnaJ_C | 0.0/**1.0**/0.0 | DnaJ;DnaJ_C (**1.0**) |
| DnaJ-1 | Q24133 | DnaJ - DnaJ_C | 0.0/**1.0**/0.0 | DnaJ;DnaJ_C (**0.99**) |
| Shv | Q9VPQ2 | DnaJ - DnaJ_C - pseudoZFLR | 0.0/0.01/**0.99** | Other (**1.0**) |
| **Class C** | | | | |
| BcDNA.GH03108 | Q9Y155 | DnaJ - zf-C2H2_jaz | 0.0/0.0/**1.0** | DnaJ;zf-C2H2_jaz (**0.99**) |
| BcDNA:GH03108 | Q9W0X8 | DnaJ - zf-C2H2_jaz | 0.0/0.0/**1.0** | DnaJ;zf-C2H2_jaz (**1.0**) |
| CG13776 | Q4V3I1 | DnaJ - DJC28_CD | 0.0/0.0/**1.0** | Other (**1.0**) |
| CG14650 | Q961F2 | DnaJ - Jiv90 | 0.0/0.0/**1.0** | Other (**1.0**) |
| CG6693 | Q9VGR7 | DnaJ | 0.0/0.0/**1.0** | Other (**1.0**) |
| CG7872 | Q9VXT2 | DnaJ | 0.0/0.0/**1.0** | Other (**0.71**) |
| CG8476 | Q8T8S9 | DnaJ | 0.0/0.0/**1.0** | Other (**0.94**) |
| Csp | Q03751 | DnaJ | 0.02/0.01/**0.97** | Other (**0.94**) |
| Dmel\CG10375 | Q9VCH9 | DnaJ | 0.0/0.0/**1.0** | Other (**0.91**) |
| Dmel\CG10565 | Q9VP77 | DnaJ - RAC_head - Myb_DNA-binding | 0.0/0.0/**1.0** | Other (**1.0**) |
| Dmel\CG14650 | Q9VN28 | DnaJ - Jiv90 | 0.0/0.0/**1.0** | Other (**1.0**) |
| Dmel\CG17187 | Q9VGR0 | DnaJ | 0.0/0.0/**1.0** | Other (**1.0**) |
| Dmel\CG30156 | Q5U0V4 | DnaJ - DUF1977 | 0.0/0.0/**1.0** | DnaJ;DUF1977 (**1.0**) |
| Dmel\CG3061 | Q9VFP0 | DnaJ - DUF1977 | 0.0/0.0/**1.0** | DnaJ;DUF1977 (**1.0**) |
| Dmel\CG32640 | Q8I079 | DnaJ | 0.0/0.0/**1.0** | Other (**1.0**) |
| Dmel\CG43322 | Q9VM54 | DnaJ - DJC28_CD | 0.0/0.0/**1.0** | Other (**1.0**) |
| Dmel\CG7130 | Q9VNW0 | DnaJ | 0.0/0.0/**1.0** | Other (**0.56**) |
| Dmel\CG7133 | Q9VNW1 | DnaJ | 0.0/0.0/**1.0** | DnaJ;DnaJ_C (**0.49**) |
| Dmel\CG7556 | Q9VWL4 | DnaJ - Myb_DNA-bind_6 | 0.0/0.0/**1.0** | Other (**1.0**) |
| Dmel\CG8476 | Q9VFY1 | DnaJ | 0.0/0.0/**1.0** | Other (**0.94**) |
| Dmel\CG8531 | Q7K0W1 | DnaJ - DnaJ-like_C11_C | 0.0/0.0/**1.0** | DnaJ;DnaJ-like_C11_C (**1.0**) |
| DnaJ-60 | P92029 | DnaJ | 0.0/0.0/**1.0** | Other (**0.98**) |
| Dph4 | Q9VNE8 | DnaJ - zf-CSL | 0.0/0.0/**1.0** | DnaJ;zf-CSL (**1.0**) |
| Hsc20 | A8JNT7 | DnaJ - HSCB_C | 0.0/0.0/**1.0** | DnaJ;HSCB_C (**1.0**) |
| NA | Q8IHG8 | DnaJ - DUF1977 | 0.0/0.0/**1.0** | DnaJ;DUF1977 (**1.0**) |
| P58IPK | Q9VHA8 | DnaJ - TPR | 0.0/0.01/**0.99** | DnaJ;TPR (**1.0**) |
| Rme-8 | A1Z7S0 | RME-8_N - GYF_2 - DnaJ | 0.0/0.0/**1.0** | Other (**1.0**) |
| Sec63 | Q9VS57 | DnaJ - Sec63 - Sec63 | 0.0/0.0/**1.0** | Other (**0.88**) |
| Tpr2 | Q9V3W1 | DnaJ - TPR | 0.0/0.0/**1.0** | DnaJ;TPR (**1.0**) |
| anon-WO0118547.426 | Q9VHW5 | DnaJ | 0.0/0.0/**1.0** | DnaJ;DnaJ_C (**0.72**) |
| Auxillin | Q9VMY8 | Pkinase - PTEN_C2 - DnaJ | 0.0/0.0/**1.0** | Other (**1.0**) |
| Jdp | Q9TVP3 | DnaJ | 0.0/0.0/**1.0** | DnaJ;DnaJ-like_C11_C (**0.74**) |
| jdp-RB | H0RNN1 | DnaJ | 0.0/0.0/**1.0** | DnaJ;DnaJ-like_C11_C (**0.74**) |
| l(3)80Fg | Q7PLG1 | DnaJ - Thioredoxin | 0.0/0.0/**1.0** | Other (**1.0**) |
| Mrj | Q7JUZ6 | DnaJ | 0.0/0.0/**1.0** | DnaJ;DnaJ_C (**0.41**) |
| Wus | Q9VX95 | TM2 - DnaJ | 0.0/0.0/**1.0** | Other (**1.0**) |

**Supplementary Table 7. Classification of the 6 *E. coli* (UP000000625 Escherichia coli (strain K12) (K12 / MG1655 / ATCC 47076)) JDPs.**

| Gene name | Uniprot | Domains | A/B/C Probabilities | Arch12 Probabilities |
|---|---|---|---|---|
| **Class A** | | | | |
| dnaJ | P08622 | DnaJ - DnaJ_C - DnaJ_CXXCXGXG | **1.0**/0.0/0.0 | DnaJ;DnaJ_C;DnaJ_CXXCXGXG (**1.0**) |
| **Canonical Class B** | | | | |
| cbpA | P36659 | DnaJ - DnaJ_C | 0.0/**1.0**/0.0 | DnaJ;DnaJ_C (**1.0**) |
| **Class C** | | | | |
| djlA | P31680 | TerB - DnaJ | 0.0/0.0/**1.0** | TerB;DnaJ (**1.0**) |
| djlB | P77381 | DnaJ; | 0.0/0.0/**1.0** | Other (**1.0**) |
| djlC | P77359 | DnaJ; | 0.0/0.0/**1.0** | Other (**1.0**) |
| hscb | P0A6L9 | DnaJ;HSCB_C; | 0.0/0.0/**1.0** | DnaJ;HSCB_C (**0.98**) |

**Supplementary Table 8. Classification of the 46 *M. mulatta* (UP000006718 Macaca mulatta (Rhesus macaque) (17573)) JDPs.**

| Gene name | Uniprot | Domains | A/B/C Probabilities | Arch12 Probabilities |
|---|---|---|---|---|
| **Class A** | | | | |
| DNAJA1 | F7C2X2 | DnaJ - DnaJ_C - DnaJ_CXXCXGXG | **0.94**/0.06/0.0 | DnaJ;DnaJ_C;DnaJ_CXXCXGXG (**0.98**) |
| DNAJA2 | G7NP88 | DnaJ - DnaJ_C - DnaJ_CXXCXGXG | **1.0**/0.0/0.0 | DnaJ;DnaJ_C;DnaJ_CXXCXGXG (**1.0**) |
| DNAJA3 | F7HGP5 | DnaJ - DnaJ_C - DnaJ_CXXCXGXG | **0.99**/0.0/0.01 | DnaJ;DnaJ_C;DnaJ_CXXCXGXG (**0.95**) |
| DNAJA4 | F7DDU9 | DnaJ - DnaJ_C - DnaJ_CXXCXGXG | **0.93**/0.06/0.01 | DnaJ;DnaJ_C;DnaJ_CXXCXGXG (**0.99**) |
| **Canonical Class B** | | | | |
| DNAJB1 | H9FTU6 | DnaJ - DnaJ_C | 0.0/**1.0**/0.0 | DnaJ;DnaJ_C (**0.98**) |
| DNAJB4 | H9EQ05 | DnaJ - DnaJ_C | 0.0/**1.0**/0.0 | DnaJ;DnaJ_C (**1.0**) |
| DNAJB5 | G7NFF1 | DnaJ - DnaJ_C | 0.0/**0.99**/0.01 | DnaJ;DnaJ_C (**0.98**) |
| DNAJB11 | G7MJ31 | DnaJ - DnaJ_C - pseudoZFLR | 0.0/0.04/**0.96** | Other (**0.89**) |
| DNAJB13 | G7NEA8 | DnaJ;DnaJ_C | 0.0/0.05/**0.95** | Other (**0.55**) |
| **Class C** | | | | |
| DNAJB12 | H9H3E3 | DnaJ - DUF1977 | 0.03/0.01/**0.97** | DnaJ;DUF1977 (**1.0**) |
| DNAJB14 | H9EUC0 | DnaJ - DUF1977 | 0.0/0.0/**1.0** | DnaJ;DUF1977 (**1.0**) |
| DNAJB2 | A0A5F7ZVN9 | DnaJ | 0.0/0.03/**0.96** | DnaJ;DnaJ_C (**0.92**) |
| DNAJB6 | A0A5F8A1X7 | DnaJ | 0.06/0.1/**0.84** | DnaJ;DnaJ_C (**1.0**) |
| DNAJB7 | A0A5F8A4E2 | DnaJ | **0.75**/0.13/0.12 | DnaJ;DnaJ_C;DnaJ_CXXCXGXG (**0.82**) |
| DNAJB9 | F7H756 | DnaJ | 0.12/0.0/**0.88** | DnaJ;DnaJ_C;DnaJ_CXXCXGXG (**0.82**) |
| DNAJC1 | F7GZD6 | DnaJ - Myb_DNA-binding | 0.0/0.0/**1.0** | Other (**1.0**) |
| DNAJC10 | A0A1D5Q0W8 | DnaJ - Thioredoxin - Thioredoxin - Thioredoxin - Thioredoxin | 0.0/0.0/**1.0** | Other (**0.99**) |
| DNAJC11 | F6RU42 | DnaJ - DnaJ-like_C11_C | 0.0/0.0/**1.0** | DnaJ;DnaJ-like_C11_C (**1.0**) |
| DNAJC12 | F6R8I6 | DnaJ | 0.0/0.0/**1.0** | Other (**1.0**) |
| DNAJC13 | A0A5F7ZCF1 | RME-8_N - GYF_2 - DnaJ | 0.0/0.0/**1.0** | Other (**1.0**) |
| DNAJC14 | A0A5F7ZGD8 | DnaJ - Jiv90 | 0.0/0.0/**1.0** | Other (**1.0**) |
| DNAJC16 | H9EUU2 | DnaJ - Thioredoxin | 0.05/0.03/**0.92** | Other (**0.61**) |
| DNAJC17 | A0A1D5RFK2 | DnaJ - RRM_1 | 0.0/0.0/**1.0** | Other (**1.0**) |
| DNAJC18 | F7CZ44 | DnaJ - DUF1977 | 0.0/0.0/**1.0** | DnaJ;DUF1977 (**1.0**) |
| DNAJC19 | F7H8X3 | DnaJ | 0.0/0.0/**1.0** | Other (**1.0**) |
| DNAJC2 | G7MM22 | DnaJ - RAC_head - Myb_DNA-binding - Myb_DNA-binding | 0.0/0.0/**1.0** | Other (**1.0**) |
| DNAJC21 | A0A1D5QQV2 | DnaJ - zf-C2H2_jaz | 0.0/0.0/**1.0** | DnaJ;zf-C2H2_jaz (**1.0**) |
| DNAJC22 | F7GW88 | TM2 - DnaJ | 0.0/0.0/**1.0** | Other (**1.0**) |
| DNAJC24 | A0A5F8ACT1 | DnaJ - PAXNEB | 0.0/0.0/**1.0** | DnaJ;zf-CSL (**1.0**) |
| DNAJC27 | G7N9J7 | Ras - DnaJ | 0.0/0.0/**1.0** | Other (**1.0**) |
| DNAJC28 | F7DMN9 | DnaJ - DJC28_CD | 0.0/0.0/**1.0** | Other (**1.0**) |
| DNAJC3 | A0A1D5Q2M0 | DnaJ - TPR | 0.0/0.0/**1.0** | DnaJ;TPR (**1.0**) |
| DNAJC30 | A0A5S6RBD8 | DnaJ | 0.0/0.0/**1.0** | DnaJ;DnaJ-X (**1.0**) |
| DNAJC4 | F6RJM6 | DnaJ | 0.0/0.0/**1.0** | Other (**0.99**) |
| DNAJC5 | A0A5F8AQ65 | DnaJ | 0.0/0.01/**0.99** | Other (**0.97**) |
| DNAJC5B | F7E930 | DnaJ | 0.0/0.0/**1.0** | Other (**0.6**) |
| DNAJC5G | F7B8I1 | DnaJ | 0.0/0.0/**1.0** | Other (**1.0**) |
| DNAJC6 | F6Y139 | PTEN_C2 - DnaJ | 0.0/0.0/**1.0** | Other (**1.0**) |
| DNAJC7 | I0FKI8 | DnaJ - TPR | 0.02/0.03/**0.95** | DnaJ;TPR (**1.0**) |
| DNAJC8 | F7HJB1 | DnaJ | 0.0/0.0/**1.0** | Other (**1.0**) |
| DNAJC9 | F6WGL3 | DnaJ | 0.0/0.0/**1.0** | Other (**1.0**) |
| GNG10 | A0A663DBJ5 | DnaJ | 0.0/0.0/**1.0** | Other (**0.95**) |
| N/A | A0A5F7ZF74 | DnaJ | 0.0/0.0/**1.0** | Other (**1.0**) |
| N/A | A0A1D5RH00 | DnaJ - TPR | 0.03/0.01/**0.96** | DnaJ;TPR (**1.0**) |
| N/A | A0A5F8AJ51 | DnaJ - Jiv90 | 0.0/0.0/**1.0** | Other (**0.94**) |
| SEC63 | F7FPB6 | DnaJ - Sec63 - Sec63 | 0.0/0.0/**1.0** | DnaJ;Sec63 (**0.97**) |

**Supplementary Table 9. Classification of the 47 *M. musculus* (UP000000589 Mus musculus (Mouse) (C57BL/6J)) JDPs.**

| Gene name | Uniprot | Domains | A/B/C Probabilities | Arch12 Probabilities |
|---|---|---|---|---|
| **Class A** | | | | |
| Dnaja1 | P63037 | DnaJ - DnaJ_C - DnaJ_CXXCXGXG | **0.78**/0.21/0.01 | DnaJ;DnaJ_C;DnaJ_CXXCXGXG (**0.99**) |
| Dnaja2 | Q9QYJ0 | DnaJ - DnaJ_C - DnaJ_CXXCXGXG | **1.0**/0.0/0.0 | DnaJ;DnaJ_C;DnaJ_CXXCXGXG (**1.0**) |
| Dnaja3 | Q99M87 | DnaJ - DnaJ_C - DnaJ_CXXCXGXG | **0.99**/0.0/0.01 | DnaJ;DnaJ_C;DnaJ_CXXCXGXG (**0.97**) |
| Dnaja4 | Q9JMC3 | DnaJ - DnaJ_C - DnaJ_CXXCXGXG | **0.93**/0.06/0.01 | DnaJ;DnaJ_C;DnaJ_CXXCXGXG (**0.99**) |
| **Canonical Class B** | | | | |
| Dnajb1 | Q9QYJ3 | DnaJ - DnaJ_C | 0.0/**1.0**/0.0 | DnaJ;DnaJ_C (**0.99**) |
| Dnajb4 | Q9D832 | DnaJ - DnaJ_C | 0.0/**0.99**/0.01 | DnaJ;DnaJ_C (**1.0**) |
| Dnajb5 | O89114 | DnaJ - DnaJ_C | 0.0/**1.0**/0.0 | DnaJ;DnaJ_C (**0.95**) |
| Dnajb11 | Q99KV1 | DnaJ - DnaJ_C - pseudoZFLR | 0.0/0.04/**0.96** | Other (**0.89**) |
| Dnajb13 | Q80Y75 | DnaJ - DnaJ_C | 0.0/0.96/ 0.04 | DnaJ;DnaJ_C (**0.96**) |
| **Class C** | | | | |
| 4930503B20Rik | Q80ZP0 | DnaJ | 0.04/0.04/**0.92** | DnaJ;DnaJ_C (**0.57**) |
| Dnajb12 | Q9QYI4 | DnaJ - DUF1977 | 0.03/0.0/**0.96** | DnaJ;DUF1977 (**1.0**) |
| Dnajb14 | Q149L6 | DnaJ - DUF1977 | 0.0/0.0/**1.0** | DnaJ;DUF1977 (**1.0**) |
| Dnajb2 | Q9QYI5 | DnaJ | 0.0/0.09/**0.91** | DnaJ;DnaJ_C (**0.88**) |
| Dnajb3 | O35723 | DnaJ | 0.01/0.39/**0.61** | DnaJ;DnaJ_C (**0.87**) |
| Dnajb6 | O54946 | DnaJ | 0.39/0.08/**0.53** | DnaJ;DnaJ_C;DnaJ_CXXCXGXG (**0.63**) |
| Dnajb7 | Q9QYI8 | DnaJ | **0.49**/0.05/0.47 | DnaJ;DnaJ_C (**0.74**) |
| Dnajb8 | Q9QYI7 | DnaJ | 0.01/0.21/**0.78** | DnaJ;DnaJ_C (**0.56**) |
| Dnajb9 | Q9QYI6 | DnaJ | 0.03/0.0/**0.97** | DnaJ;DnaJ_C;DnaJ_CXXCXGXG (**0.66**) |
| Dnajc1 | Q61712 | DnaJ - Myb_DNA-binding | 0.0/0.0/**1.0** | Other (**1.0**) |
| Dnajc10 | Q9DC23 | DnaJ - Thioredoxin - Thioredoxin - Thioredoxin - Thioredoxin | 0.0/0.0/**1.0** | Other (**0.97**) |
| Dnajc11 | Q5U458 | DnaJ - DnaJ-like_C11_C | 0.0/0.0/**1.0** | DnaJ;DnaJ-like_C11_C (**1.0**) |
| Dnajc12 | Q9R022 | DnaJ | 0.0/0.0/**1.0** | Other (**1.0**) |
| Dnajc13 | D4AFX7 | RME-8_N - GYF_2 - DnaJ | 0.0/0.0/**1.0** | Other (**1.0**) |
| Dnajc14 | Q921R4 | DnaJ - Jiv90 | 0.0/0.0/**1.0** | Other (**1.0**) |
| Dnajc16 | Q80TN4 | DnaJ - Thioredoxin | 0.0/0.05/**0.95** | Other (**0.79**) |
| Dnajc17 | Q91WT4 | DnaJ - RRM_1 | 0.0/0.0/**1.0** | Other (**1.0**) |
| Dnajc18 | Q9CZJ9 | DnaJ - DUF1977 | 0.0/0.0/**1.0** | DnaJ;DUF1977 (**1.0**) |
| Dnajc19 | Q9CQV7 | DnaJ | 0.0/0.0/**1.0** | Other (**1.0**) |
| Dnajc2 | P54103 | DnaJ - RAC_head - Myb_DNA-binding - Myb_DNA-binding | 0.0/0.0/**1.0** | Other (**1.0**) |
| Dnajc21 | E9Q8D0 | DnaJ - zf-C2H2_jaz | 0.0/0.0/**1.0** | DnaJ;zf-C2H2_jaz (**1.0**) |
| Dnajc22 | Q8CHS2 | TM2 - DnaJ | 0.0/0.0/**1.0** | Other (**1.0**) |
| Dnajc24 | Q91ZF0 | DnaJ - zf-CSL | 0.0/0.0/**1.0** | DnaJ;zf-CSL (**1.0**) |
| Dnajc25 | A2ALW5 | DnaJ | 0.0/0.0/**1.0** | Other (**0.88**) |
| Dnajc27 | Q8CFP6 | Ras - DnaJ | 0.0/0.0/**1.0** | Other (**1.0**) |
| Dnajc28 | Q8VCE1 | DnaJ - DJC28_CD | 0.0/0.0/**1.0** | Other (**1.0**) |
| Dnajc3 | Q91YW3 | DnaJ - TPR | 0.0/0.0/**1.0** | DnaJ;TPR (**1.0**) |
| Dnajc30 | P59041 | DnaJ | 0.0/0.0/**1.0** | DnaJ;DnaJ-X (**1.0**) |
| Dnajc4 | Q9D844 | DnaJ | 0.0/0.0/**1.0** | Other (**0.74**) |
| Dnajc5 | P60904 | DnaJ | 0.0/0.01/**0.99** | Other (**0.97**) |
| Dnajc5b | Q9CQ94 | DnaJ | 0.0/0.0/**0.99** | Other (**0.91**) |
| Dnajc5g | Q8C632 | DnaJ | 0.0/0.0/**1.0** | Other (**0.99**) |
| Dnajc7 | Q9QYI3 | DnaJ - TPR | 0.02/0.03/**0.95** | DnaJ;TPR (**1.0**) |
| Dnajc8 | Q6NZB0 | DnaJ | 0.0/0.0/**1.0** | Other (**1.0**) |
| Dnajc9 | Q91WN1 | DnaJ | 0.0/0.0/**1.0** | Other (**0.95**) |
| Gak | Q99KY4 | Pkinase - PTEN_C2 - DnaJ | 0.0/0.0/**1.0** | Other (**1.0**) |
| Gm20503 | G3UZK1 | DnaJ - G-gamma | 0.0/0.0/**1.0** | Other (**0.96**) |
| Sec63 | Q8VHE0 | DnaJ - Sec63 | 0.0/0.0/**1.0** | DnaJ;Sec63 (**0.97**) |

### Supplementary Table 10. Classification of the 46 *R. norvegicus* (UP000002494 Rattus norvegicus (Rat) (Brown Norway)) JDPs.

| Gene | Entry | Domains | A/B/C Probabilities | Arch12 Probabilities |
|---|---|---|---|---|
| **Class A** | | | | |
| Dnaja1 | P63036 | DnaJ - DnaJ_C - DnaJ_CXXCXGXG | **0.78**/0.21/0.01 | DnaJ;DnaJ_C;DnaJ_CXXCXGXG (**0.99**) |
| Dnaja2 | O35824 | DnaJ - DnaJ_C - DnaJ_CXXCXGXG | **1.0**/0.0/0.0 | DnaJ;DnaJ_C;DnaJ_CXXCXGXG (**1.0**) |
| Dnaja3 | A0A0G2K5E4 | DnaJ - DnaJ_C - DnaJ_CXXCXGXG | **0.99**/0.0/0.01 | DnaJ;DnaJ_C;DnaJ_CXXCXGXG (**0.97**) |
| Dnaja4 | Q4QR73 | DnaJ - DnaJ_C - DnaJ_CXXCXGXG | **0.93**/0.06/0.01 | DnaJ;DnaJ_C;DnaJ_CXXCXGXG (**0.99**) |
| **Canonical Class B** | | | | |
| Dnajb1 | B0K030 | DnaJ - DnaJ_C | 0.0/**1.0**/0.0 | DnaJ;DnaJ_C (**0.99**) |
| Dnajb11 | Q6TUG0 | DnaJ - DnaJ_C - pseudoZFLR | 0.0/0.04/**0.96** | Other (**0.89**) |
| Dnajb4 | Q5XIP0 | DnaJ - DnaJ_C | 0.0/**0.99**/0.01 | DnaJ;DnaJ_C (**1.0**) |
| Dnajb5 | D3ZB76 | DnaJ - DnaJ_C | 0.0/**1.0**/0.0 | DnaJ;DnaJ_C (**0.95**) |
| Dnajb13 | Q5YKV6 | DnaJ – DnaJ_C | 0.0/0.46/**0.54** | DnaJ;DnaJ_C (**0.77**) |
| **Class C** | | | | |
| Dnajb12 | Q5FVC4 | DnaJ - DUF1977 | 0.03/0.0/**0.96** | DnaJ;DUF1977 (**1.0**) |
| Dnajb14 | A0A0G2JTM9 | DnaJ - DUF1977 | 0.0/0.0/**1.0** | DnaJ;DUF1977 (**1.0**) |
| Dnajb2 | D4ABC2 | DnaJ | 0.0/0.0/**1.0** | DnaJ;TPR (**0.9**) |
| Dnajb3 | D3ZAC5 | DnaJ | 0.0/**0.87**/0.12 | DnaJ;DnaJ_C (**0.64**) |
| Dnajb6 | Q6AYU3 | DnaJ | 0.39/0.08/**0.53** | DnaJ;DnaJ_C;DnaJ_CXXCXGXG (**0.63**) |
| Dnajb7 | D3ZN46 | DnaJ | **0.63**/0.05/0.31 | DnaJ;DnaJ_C;DnaJ_CXXCXGXG (**0.83**) |
| Dnajb8 | D3ZWI8 | DnaJ | 0.01/0.21/**0.78** | DnaJ;DnaJ_C (**0.56**) |
| Dnajb9 | P97554 | DnaJ | 0.07/0.0/**0.92** | DnaJ;DnaJ_C;DnaJ_CXXCXGXG (**0.88**) |
| Dnajc1 | F1LVX1 | DnaJ - Myb_DNA-binding | 0.0/0.0/**1.0** | Other (**1.0**) |
| Dnajc10 | Q498R3 | DnaJ - Thioredoxin - Thioredoxin - Thioredoxin - Thioredoxin | 0.0/0.0/**1.0** | Other (**0.97**) |
| Dnajc11 | B1WBY5 | DnaJ - DnaJ-like_C11_C | 0.0/0.0/**1.0** | DnaJ;DnaJ-like_C11_C (**1.0**) |
| Dnajc12 | Q925T0 | DnaJ | 0.0/0.0/**1.0** | Other (**1.0**) |
| Dnajc13 | D3ZN27 | RME-8_N - GYF_2 - DnaJ | 0.0/0.0/**1.0** | Other (**1.0**) |
| Dnajc14 | Q5XIX0 | DnaJ - Jiv90 | 0.0/0.0/**1.0** | Other (**1.0**) |
| Dnajc16 | Q5FVM7 | DnaJ - Thioredoxin | 0.0/0.04/**0.96** | Other (**0.61**) |
| Dnajc17 | D3ZSC8 | DnaJ - RRM_1 | 0.0/0.0/**1.0** | Other (**1.0**) |
| Dnajc18 | Q6AXW3 | DnaJ - DUF1977 | 0.0/0.0/**1.0** | DnaJ;DUF1977 (**1.0**) |
| Dnajc2 | Q7TQ20 | DnaJ - RAC_head - Myb_DNA-binding - Myb_DNA-binding | 0.0/0.0/**1.0** | Other (**1.0**) |
| Dnajc21 | A0A0G2QC19 | DnaJ - zf-C2H2_jaz | 0.0/0.0/**1.0** | DnaJ;zf-C2H2_jaz (**1.0**) |
| Dnajc22 | Q5PR00 | TM2 - DnaJ | 0.0/0.0/**1.0** | Other (**1.0**) |
| Dnajc24 | F1M2S2 | DnaJ - zf-CSL | 0.0/0.0/**1.0** | DnaJ;zf-CSL (**1.0**) |
| Dnajc25 | Q5BJW9 | DnaJ | 0.0/0.0/**1.0** | Other (**0.88**) |
| Dnajc27 | Q6IML7 | Ras - DnaJ | 0.0/0.0/**1.0** | Other (**1.0**) |
| Dnajc28 | Q6AYG2 | DnaJ - DJC28_CD | 0.0/0.0/**1.0** | Other (**1.0**) |
| Dnajc3 | Q9R0T3 | DnaJ - TPR | 0.0/0.0/**1.0** | DnaJ;TPR (**1.0**) |
| Dnajc30 | B2RYV0 | DnaJ | 0.0/0.0/**1.0** | DnaJ;DnaJ-X (**1.0**) |
| Dnajc4 | Q5M867 | DnaJ | 0.0/0.0/**1.0** | Other (**0.49**) |
| Dnajc5 | P60905 | DnaJ | 0.0/0.01/**0.99** | Other (**0.97**) |
| Dnajc5b | D3ZD82 | DnaJ | 0.0/0.0/**1.0** | Other (**0.48**) |
| Dnajc5g | Q5XIK9 | DnaJ | 0.0/0.11/**0.89** | Other (**0.72**) |
| Dnajc7 | A0A0G2K435 | DnaJ - TPR | 0.02/0.03/**0.95** | DnaJ;TPR (**1.0**) |
| Dnajc8 | Q642C0 | DnaJ | 0.0/0.0/**1.0** | Other (**1.0**) |
| Dnajc9 | A8QIC3 | DnaJ | 0.0/0.0/**1.0** | Other (**0.85**) |
| Gak | P97874 | Pkinase - PTEN_C2 - DnaJ | 0.0/0.0/**1.0** | Other (**1.0**) |
| Samd13 | Q9QZW8 | DnaJ | 0.0/0.0/**0.99** | DnaJ;DnaJ_C;DnaJ_CXXCXGXG (**0.49**) |
| Sec63 | A0A0G2JUJ9 | DnaJ - Sec63 - Sec63 | 0.0/0.0/**1.0** | DnaJ;Sec63 (**0.97**) |
| Zrf4 | Q7TQ19 | DnaJ | 0.0/0.0/**1.0** | Other (**1.0**) |
